# KATMAP: Inferring splicing factor activity and regulatory targets from knockdown data

**DOI:** 10.1101/2024.06.25.600605

**Authors:** Michael P. McGurk, David C. McWatters, Christopher B. Burge

## Abstract

Typical RNAseq experiments uncover hundreds of splicing changes, reflecting underlying changes in splicing factor (SF) activity. Understanding transcriptomic variation in terms of SF activity requires elucidating the rules by which each SF impacts splicing. Here we present an interpretable regression model, KATMAP, which models splicing changes transcriptome-wide in terms of changes in SF binding and resulting altered regulation. The regulatory principles KATMAP learns generalize to predict the SF’s regulation at individual exons, with potential for design of splice-switching antisense oligonucleotides and inference of the displaced factor. We also discover cooperative splicing regulation by QKI and RFBOX proteins. KATMAP interprets RNAseq data by uncovering the factors responsible for transcriptomic changes, distinguishing direct SF targets from indirect effect, and infers relevant SFs from clinical RNAseq data.

## Main

Programs of transcriptional and post-transcriptional regulation are responsible for much of development, physiology and disease. Orchestrating these gene expression programs are regulatory factors which recognize sequence elements distributed across the genome and transcriptome. The transcriptomic differences between two biological states are typically large, with hundreds or thousands of genes differing in splicing or expression, but these differences typically result from a much smaller set of regulatory factors. Disentangling gene expression programs in terms of regulatory factors and their targets is critical to understanding the genomic basis of physiology, and doing so requires determining the rules by which these factors exert their activities.

Alternative splicing greatly expands proteomic diversity by allowing a single gene to generate dozens of distinct splicing isoforms (spliceoforms), often with functions tuned to specific tissues or developmental stages. Between 10–20% of disease-causing mutations are estimated to result in changes to splicing levels of individual exons (Jaganathan et al. 2019; Lim et al. 2011) while perturbations of splicing factor (SF) genes can broadly disrupt splicing. Several SFs are known oncogenes or tumor suppressors (Bradley and Anczuków 2023; Karni et al. 2007) and the resulting disruptions to splicing programs represent vulnerabilities that can be exploited (North et al. 2022). Splicing defects at individual exons can often be corrected by splice-switching antisense oligonucleotides (ssASOs) (Finkel et al. 2016; Hua et al. 2007). However, brute force tiling is often required to identify effective ASOs because the relevant SFs and splicing regulatory elements (SREs) are unknown. Interpretation of clinical variants that may impact splicing is also made challenging by our incomplete knowledge of SF activities and SREs (Ellingford et al. 2022). Elucidating precise rules governing SF activity would facilitate ASO design, variant interpretation, and the identification of SFs that contribute to disease.

Despite the recent popularity of deep learning models (Jaganathan et al. 2019), there is a fundamental limitation in the application of black box “AI” models to the study of gene regulation. The way in which a pre-mRNA is processed is not an intrinsic property of its sequence, but depends on the cellular context in which the processing reactions occur. Black box models do not formulate their predictions in terms of biological hypotheses and so do not offer a means to separate the context-independent (e.g. the regulatory activity and binding affinities of regulatory factors) from the context-dependent aspects of regulation (e.g. the expression level of regulatory factors), limiting the extent to which they can be meaningfully generalized beyond their training set. Bayesian generative models offer a resolution by explicitly describing how data arose in terms biological parameters, providing interpretable explanations that not only generate hypothesis, but facilitate the translation of these insights to applied settings.

Partial solutions are offered by approaches that assign regulatory activity to hundreds or thousands of motifs or *k* -mers from transcriptome data without using prior knowledge of transcription (Balwierz et al. 2014; J. D. Rubin et al. 2021), stability (Krismer et al. 2020), or splicing (Bak et al. 2024; Hugh-White 2021). But these approaches do not extract generalizable insights or even directly identify the causal factors, because they require discovering the putative activity of each motif every time they are applied. More complete solutions have leveraged perturbation experiments to predict the targets of specific SFs, by incorporating expert knowledge of the factor and using crosslinking/immunoprecipitation (CLIP) data to inform binding sites (Han et al. 2014; Weyn-Vanhentenryck et al. 2014; C. Zhang et al. 2010). However, the challenge of crafting these models to the specific details of the SF’s binding and activity has limited their application to a few well-studied factors and a reliance on crosslinking data still ties the insights to exons where the SF was detectably bound in specific experimental data.

Here we present a flexible and interpretable model for learning regulatory activity from knockdown or overexpression RNAseq data, which can then be applied to generate predictions and interpret sequencing data. Our approach, Knockdown Activity and Target Models from Additive regression Predictions (KATMAP), explicitly models the effect of knockdown by assigning changes in SF occupancy at potential binding sites throughout the transcriptome to predict regulation at individual exons (Fig. 1a). We utilize a hierarchical framework that tailors modelling assumptions to each SF and learns the nonlinear biophysical relationship between motif affinity and binding, allowing KATMAP to be broadly applied off-the-shelf to diverse SF datasets. Learning a regulatory model requires as inputs only the results of a perturbation experiment and some model of the factor’s RNA sequence specificity, such as a position-weight matrix (PWM) (Fig.1b). Once learned, these models can be applied to any exon sequence to identify cis-elements and associated SFs, or to any RNAseq dataset to infer the SFs that explain splicing differences between conditions, for example to identify disease drivers (Fig. 1c).

**Figure 1:**
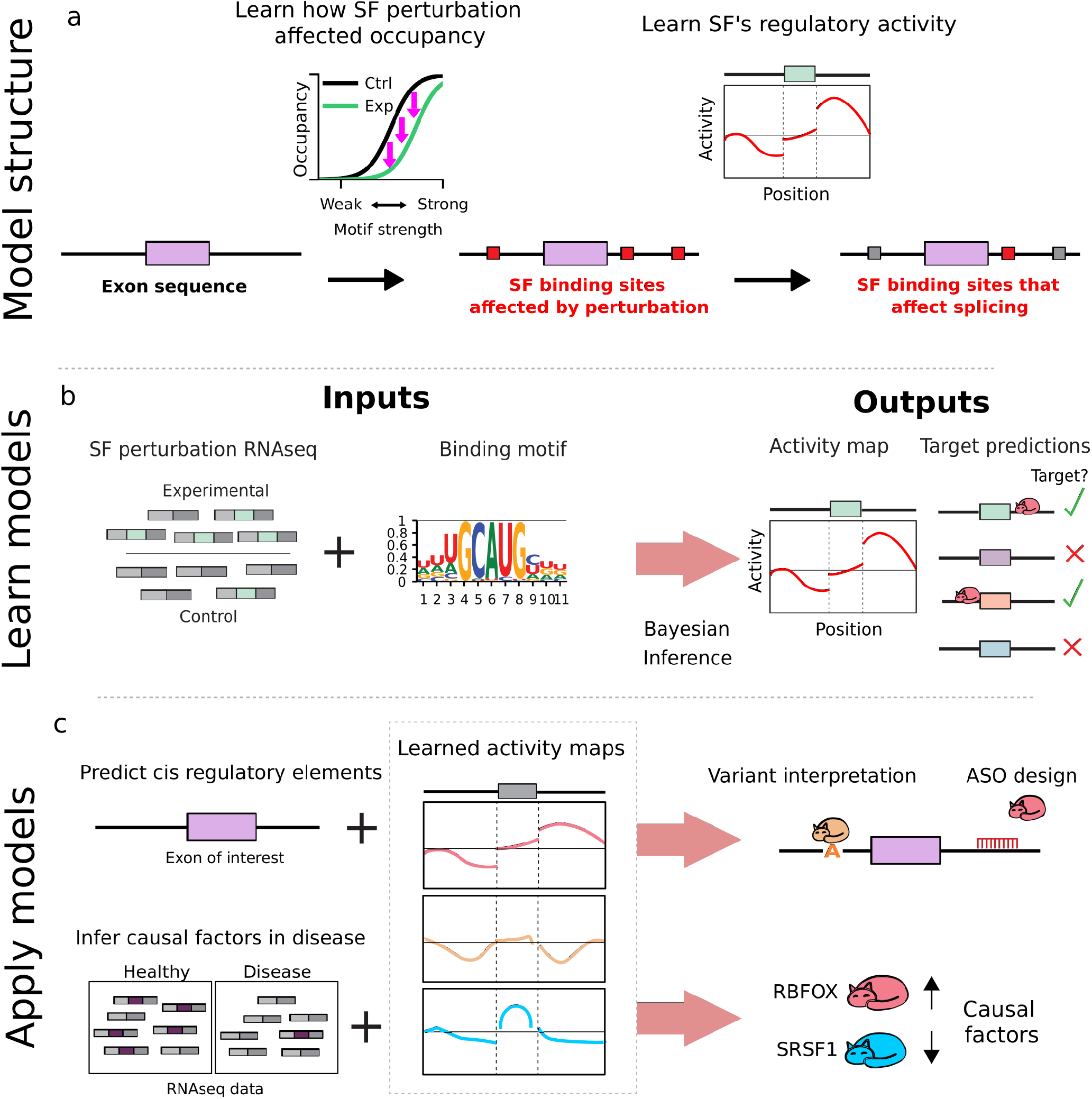
High-level overview of the KATMAP model and its applications. **a)** The general workflow of how KATMAP predicts the regulation of individual exons based on their sequence. **b)** Learning the rules by which an SF regulates splicing requires only an RNAseq dataset where that SF was perturbed and some model of its sequence specificity. Bayesian inference determines which splicing models are consistent with the observed splicing changes, yielding an activity map describing the SF’s position-specific regulatory activity and identifying which exons are teh SF’s direct targets and which splicing changes represent indirect effects. **c)** Examples of how learned models can be applied to interpret splicing in terms of SF regulation. **Top:** Scoring an exon of interest to predict the SREs and SFs that control its splicing, with application to variant interpretation and ASO design. **Bottom:** Identifying the causal SFs underlying splicing differences observed in a given RNAseq dataset.

## Results

### Overview of the KATMAP model

The intuition guiding our model is as follows. Prior to a perturbation in the level of a sequence-specific splicing factor (SF), sequence motifs in the transcriptome are bound by the SF at varying levels of occupancy. We assume the factor can only influence the splicing reaction if its binding sites are properly situated relative to the exon and that binding elsewhere has negligible effect. After knockdown, the free protein concentration decreases, leading to reduced occupancy across the protein’s binding sites. If binding is lost from a regulatory region, then loss of regulation results in a change in exon inclusion. The greater the decrease in binding, the more likely it is to affect inclusion. The logic is similar for SF overexpression, except that free protein concentration, occupancy, and regulation increase upon perturbation.

Following this logic, KATMAP models splicing changes in terms of changes in SF binding to the transcript. As inputs, our model requires only some information about a SF’s RNA sequence specificity and the results of a differential splicing analysis which has labelled each exon as upregulated, downregulated or “not significant”. The model aims to predict whether and in which direction each exon changes in splicing by answering two questions: 1) Are there binding sites near the exon which were affected by the SF knockdown? 2) Where must the SF bind near an exon to regulate its inclusion? We frame this as a generalized additive model which represents SF binding and regulation with nonlinear terms but accounts for other features predictive of detectable splicing change (such as gene expression level) through additional linear terms.

Our model begins by scoring all potential binding motifs around an exon’s splice sites, extending 200 nt on the intronic and 60 nt on the exonic side, using position weight matrices (PWMs), position-specific affinity matrices (PSAMs), or other affinity models (Supplementary fig. 1a). These *motif scores* are then transformed to changes-in-binding using a *binding function* based on a biophysical description of how occupancy changes after a decrease in SF availability (Supplementary fig. 1b, c). The shape of the binding function is controlled by a *most-relevant-affinity* parameter that defines which binding sites were most affected by the knockdown (i.e. shifts the function along the X-axis) and an *affinity scale* parameter which defines the range of motif scores affected by the knockdown (i.e. widens/narrows the function); we refer to these collectively as the *binding parameters* (see Methods).

We represent the SF’s position-specific regulatory activity with a set of parameters termed the *activity map* (Supplementary fig. 1d), which assigns enhancing (*>* 0) or suppressing (*<* 0) regulatory activity to different positions on the intronic and exonic sides of 3SS and 5SS. These regulatory activities are shared across all exons in the analysis. Critically, we assume that activity varies smoothly with position, with discontinuities at the exon-intron boundaries that allow exonic and intronic binding to have distinct activities; how smoothly depends on two *spatial correlation* parameters. We multiply each change-in-binding by the corresponding activity to determine its impact on the splicing outcome (Supplementary fig. 1e). The sum of these products defines the SF’s *splicing impact* on the exon, making the simplifying assumption that regulation is additive.

Finally, we incorporate these splicing impacts into a regression framework which assigns each exon a probability of being upregulated, downregulated or unchanged (Supplementary fig. 1f). In addition to SF binding, the unperturbed level of exon inclusion and gene expression impact our statistical power to detect splicing changes. Specifically, exons with inclusion levels (PSI values) near the boundaries of 0% or 100% inclusion are less likely to be detected as down- or upregulated, respectively, while statistical power to detect splicing changes is limited for exons in genes with low expression. We include these confounding variables (e.g. unperturbed inclusion level, expression) and their squares as linear predictors, whose effects are controlled for by a set of *linear coefficients*.

This model architecture answers the two questions we posed in an integrated fashion. SF binding and regulatory activity of the perturbed SF are modeled simultaneously, without the need to define arbitrary cutoffs for binding or activity. The activity map provides a simple description of the SF’s position-specific activity, identifying where a binding site must occur to influence an exon’s inclusion positively or negatively, an intuitive notion of the SF’s function which we propose as an alternative to the traditional RNA map. Assuming that this activity changes smoothly with position (except at exon-intron boundaries) is both biologically plausible (Graveley, Hertel, and Maniatis 1998) and improves statistical power by pooling information spatially (Supplementary fig. 1g, h). How smoothly it changes is automatically learned from the data.

The goal of KATMAP’s inference algorithm is to determine which parameter values provide good explanations for the observed splicing changes. Prior target prediction models obtained point estimates of the most likely parameter values (C. Zhang et al. 2010), but here we wished to also quantify our uncertainty by identifying the range of plausible hypotheses. We therefore used Bayesian inference to precisely evaluate the plausibility of hypotheses given observed data. We structured KATMAP’s model such that posterior inference could be split into two tractable subproblems. Loosely speaking, the parameters used to denote binding (*most-relevant-affinity* and *affinity-scale*) and spatial correlation can be considered *hyperparameters*. If the hyperparameters are held constant, the model simplifies to a generalized linear model (GLM), such that the dozens of *activity* and *linear* coefficients can be identified by straightforward optimization (Rue, Martino, and Chopin 2009). To leverage this property, we couple adaptive importance sampling with integrated nested Laplace approximations (INLA) (Rue, Martino, and Chopin 2009), dividing posterior inference into: 1) an inner step that solves conditional models to learn which activity and linear coefficients best explain the data assuming particular binding and spatial correlation parameter values to be true; and 2) an outer step that evaluates a marginal model to learn which binding and spatial correlation parameter values are consistent with the data. This approach yields samples from the joint posterior that span the range of plausible parameter values (Supplementary fig. 1i), allowing us to compute point estimates of the most plausible parameters, quantify our uncertainty with credible intervals, and propagate this uncertainty forward into our predictions of regulation.

### KATMAP accurately infers parameters and quantifies uncertainty

To validate our inference strategy, we simulated 21 SF knockdown datasets with known parameter values and found that KATMAP robustly recovers the true parameters (Supplementary fig. 2a-c). Importantly, our posterior samples properly quantify the uncertainty regarding the activity and binding parameters (Supplementary fig. 2e, f). Because the activity and binding parameters are accurately recovered, so are the splicing impacts used to simulate the splicing changes (Supplementary fig. 2d). Linear coefficients are slightly underestimated Supplementary fig. 2c, h), likely due to the multivariate Gaussian approximation to their conditional posterior (Rue, Martino, and Chopin 2009); these parameters serve primarily to control for potential confounders, and this subtle bias is unlikely to interfere with this.

To assess performance on real data, we selected ten knockdowns with many splicing changes. For each experiment, we trained on 70% of the differentially spliced exons, holding out the remaining 30% as a test set. We evaluated our models against these held-out exons, comparing the true splicing changes to our predictions of which exons were up- or down-regulated or unchanged upon knockdown. In all ten evaluations, the predicted probabilities were well-calibrated, with the probability that an exon was affected by knockdown in a given direction corresponding to the frequency of that outcome (Supplementary fig. 2h, i)

### KATMAP robustly infers splicing regulatory activity

We applied KATMAP to knockdown data from the ENCODE project for 35 SFs for which we also had good models of the protein’s sequence specificity (Supplementary Table 1) (Van Nostrand et al. 2020). We used RNA-bind-n-seq-derived position-specific affinity matrices (PSAMs) as the binding model for 23 SFs (Jens et al. 2022; Lambert et al. 2014), and RNAcompete-derived position weight matrices (PWMs) for the remaining 12 (Ray, Kazan, Chan, et al. 2009; Ray, Kazan, Cook, et al. 2013). Eighteen SFs showed significant enhancing or repressing activity maps and outperformed a reduced model (with splicing activity excluded) in at least one knockdown, with 9 displaying enhancing activity, 6 showing repressing activity, and 3 with both enhancing and repressing activity (Fig. 2a, Supplementary Data 1). Among these eighteen, 14 had available knockdown/RNA-seq data from both lymphocyte-derived K562 cells and hepatocyte-derived HepG2 cells. For all but one factor (SF1), the activity maps were consistent between the two knockdowns, highlighting the reproducibility of our inferences (Fig. 2b).

**Supplementary Figure 1:**
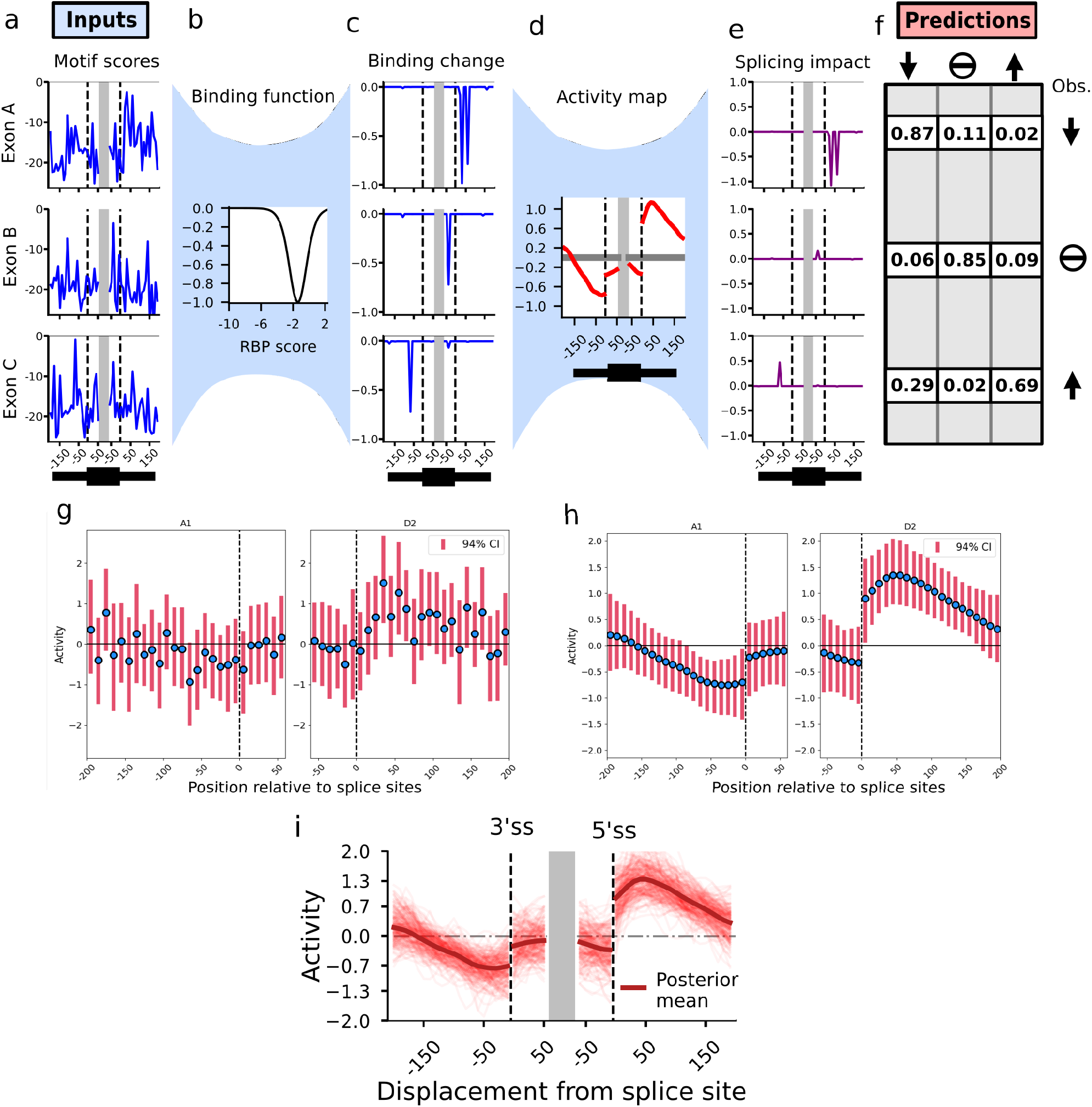
Detailed overview of model architecture. **a-f)** Examples of the model applied to three real exons in the *RBFOX2* (HepG2) knockdown. **a)** RBFOX2 motif scores assigned to sequences around the 3’ and 5’ss. **b**) The binding function that transforms the scores into relative changes in binding. **c)** Predicted changes in binding obtained by passing the motifs scores (**a**) through the binding function (**b**). **d)** An activity map describing the effect on splicing of RBFOX2 binding at positions near the 3’ and 5’ss. Discontinuities at the splice sites allow activity to differ between introns and exons. **e)** The predicted splicing impacts, computed by multiplying the binding changes (**c**) by the activity map (**d**). **f)** The predicted probabilities of being downregulated (↓), unchanged (–), or upregulated (↑) for these exons and the observed splicing changes. **g, h)** The activity maps inferred from an RBFOX2 knockdown in HepG2 cells with priors assuming independence among the activity coefficients (**g**) or with our full model’s assumption of spatial correlation (**h**). **i)** A posterior sample of 200 activity maps for the RBFOX2 (HepG2) KD.

**Supplementary Figure 2:**
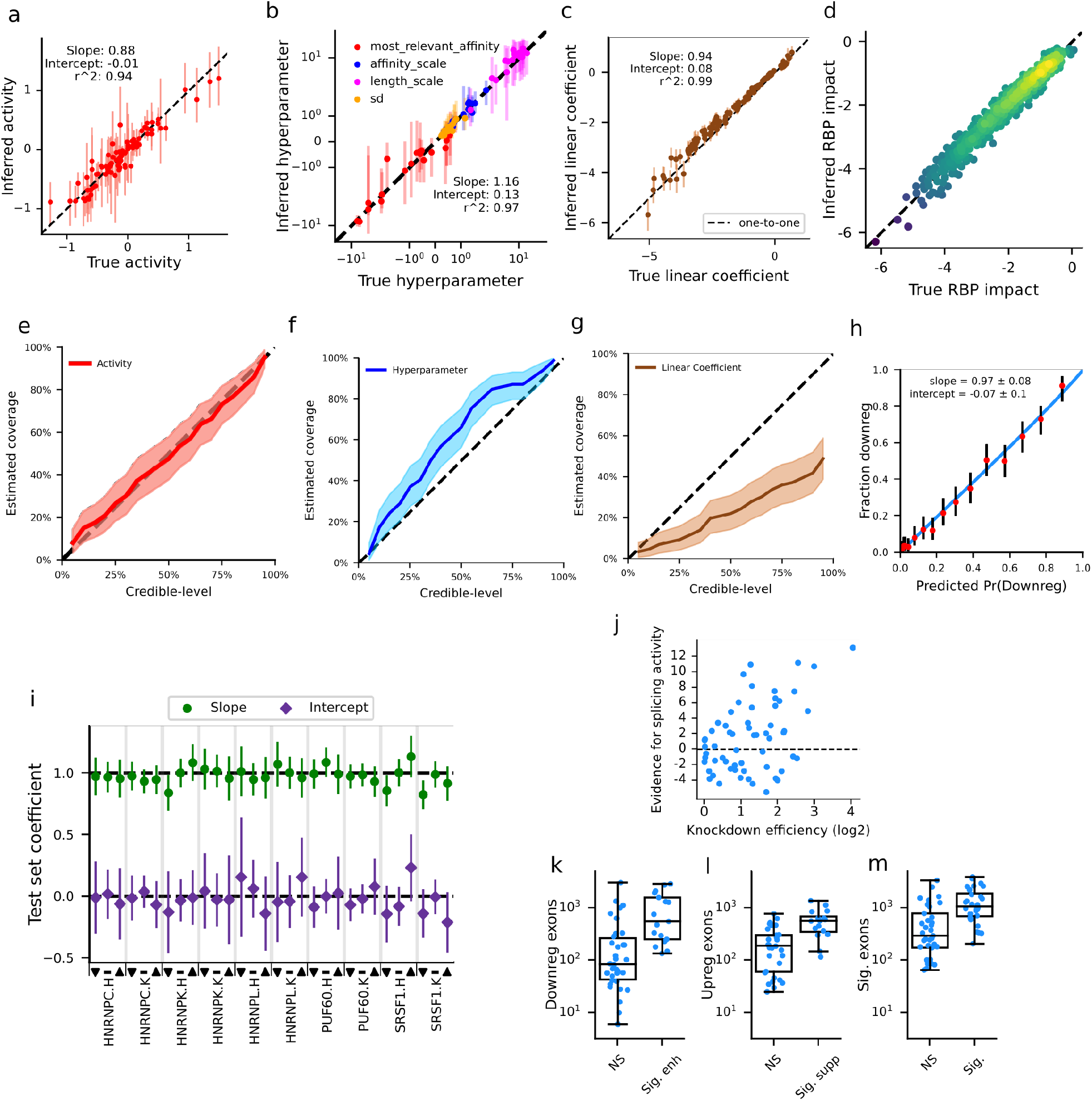
Assessment of KATMAP’s performance. **a-c)** True activities (**a**), hyperparameters (**b**), and linear coefficients (**c**) used in the 20 simulated datasets versus the values (with 94%-CI) recovered by inference. Only four activities per simulation are shown to ensure independence of the plotted values. **d)** An example comparing splicing impacts used to simulate splicing changes and the splicing impacts recovered by inference. **e-g)** Coverage plots of how frequently credible intervals contain the true activity (**e**), hyperparameter (**f**), or linear coefficient (**g**), for different CI-levels. **h)** Calibration plot for the PUF60 (K562) knockdown, comparing average *Pr*(*Downreg*) and fraction truly downregulated for sets of 200 held out exons. The blue line is a binomial regression fit, the dots are posterior means with 94%-credible intervals.**i)** The slopes and intercepts of binomial regressions evaluating whether the downreg (▽), unchanged (–), and upreg (△) probabilities correctly describe the splicing changes at held out exons, as in **h**, for ten knockdowns. The dotted lines are the expected values of the slope (1) and intercept (0) for well-calibrated models. **j)** The relationship between the efficiency of an ENCODE knockdown as shrinkage estimate of the gene’s of log2 fold-change in the DEseq analysis. **k-m)** The numbers of differentially spliced exons in knock-downs where KATMAP identified enhancing (**k**) or suppressing (**l**) activity, or activity in either direction (**m**). Boxes are upper and lower quartiles, and whiskers extend to the furthest data points from the box within 1.5 of the interquartile range (IQR).

The evidence KATMAP identified for splicing activity correlated with knockdown efficiency (*r* =0.45, Supplementary fig. 2j), with stronger knockdowns providing more evidence for splicing activity. Similarly, a sufficient number of splicing changes was required to infer activity, with evidence of enhancing or repressing activity only identified in knockdowns where at least 100 exons were down- or upregulated, respectively (Supplementary fig. 2k-m). For example, both ENCODE knockdowns for the splicing activator *TRA2A* yielded *<* 100 downregulated exons and no clear evidence of activity–perhaps because of compensation by paralog *TRA2B*. However, applying our model to a *TRA2A/B* double knockdown experiment with *>* 1000 downregulated exons (Best et al. 2014) recovered strong evidence of the expected exonic enhancing activity (Fig. 2c).

Examining the significant activity maps identifies three distinct groups of splicing activators. First, we identify SRSF1, SRSF7, and TRA2A/B as exhibiting the classic exonic enhancing pattern of SR protein regulation (Fig. 2a,c). TRA2A/B shows similar activity near both splice sites, while SRSF1 is inferred to have stronger activity near the 5SS and SRSF7 near the 3SS. Second, several polypyrimidine-binding factors show enhancing activity directly upstream of the 3SS, consistent with polyY tract definition. For U2AF2– which canonically defines the polyY tract (Merendino et al. 1999)–and its homolog PUF60 (Page-McCaw, Amonlirdviman, and Sharp 1999) we infer intronic enhancing activity just upstream of the 3SS (Fig. 2a,d). U2AF2’s inferred activity is constrained to within 20 nt of the 3SS, while PUF60’s activity is slightly more upstream and extends 40-60 nt into the intron, consistent with observations of extended polyY tracts at PUF60-responsive exons (Královičová et al. 2018). Similar upstream enhancing activity is inferred for PCBP1 (Fig. 2a), but with additional activity downstream of the 5SS, consistent with prior observations of C-rich motifs up- and downstream of PCBP1- and PCBP2-activated exons (Ji et al. 2016). KATMAP failed to detect the expected enhancing activity of the branch point-binding factor SF1 and the exonic enhancing activity of SRSF5, perhaps due to indirect effects of other SFs obscuring the signal in these knockdowns.

While upstream activators display greatest activity near the 3SS, a third group of factors with downstream enhancing activity–DAZAP1, HNRNPF/H, QKI, and RBFOX–have their greatest activity tens of nucleotides away from the 5SS and influence splicing from a wider intronic window (Fig. 2a). Among these, two pairs of factors are strikingly similar in their activity. The related poly-G binding factors HNRNPF and HNRNPH1 both enhance inclusion when bound in the downstream intron while suppressing inclusion in the exon. HNRNPH1 was one of the first SFs demonstrated to activate and repress inclusion in a position-dependent manner (C. D. Chen, Kobayashi, and Helfman 1999; Chou et al. 1999). RBFOX2 and QKI also share patterns of activity (Fig. 2e,f)–enhancing downstream while suppressing upstream–but, unlike HNRNPF/H, are not related and bind distinct motifs (YGCAUG versus ACUAAC, respectively).

In contrast to splicing activators, all inferred repressors displayed activity in multiple regions (Fig. 2a). The PCBP-related HNRNPK protein shows silencing activity upstream of the 3SS and in the exon (Fig. 2g), consistent with its role as a splicing repressor (Cao et al. 2012), and observations that PCBP2 and HNRNPK can regulate the same exon in opposing directions (Ghanem et al. 2018). We find a similar pattern of upstream and exonic silencing activity for PTBP1 (polypyrimidine tract binding protein), as well as evidence of downstream enhancing activity (Fig. 2a), all of which is consistent with prior studies (Llorian et al. 2010). This exonic activity of HNRNPK and PTBP1 stands in contrast to the pyrimidine-binding upstream activators–which lacked exonic activity. The other repressors have even wider regions of suppression. HNRNPC’s silencing activity is strongest in the ∼ 100 nt up- and downstream of the exon, but has two additional regions of silencing activity at +/- 200 nt (Fig. 2h). HNRNPC is know to form tetramers, with repression suggested to result from looping out the exon, which may explain the multiple regions of activity (Zarnack et al. 2013). HNRNPL activity maps also display repressing activity both up- and downstream (Fig. 2a), but further represses within the exon, consistent with prior work (Heiner et al. 2010; Rothrock, House, and Lynch 2005). MATR3 (Matrin3) has even broader regions of repression, extending throughout both up- and downstream introns in the HepG2 model, but only showing evidence of upstream repression in K562 cells (Fig. 2a). This broad silencing activity is consistent with CLIP-based enrichment maps of MATR3-repressed exons which found MATR3 uniformly distributed hundreds of bases into the flanking introns (Coelho et al. 2015).

The diverse position-dependent patterns of splicing regulation we uncover underscore the power and flexibility of KATMAP’s approach to describing activity. Assuming only that activity varies smoothly with distance (and learning how smoothly from the data) allowed KATMAP to detect both broad and narrow ranges of activity, within, upstream, or downstream of target exons. We were able to robustly and reproducibly uncover this activity by modelling the sequence of exons and without relying on crosslinking data, avoiding the bias toward exonic signal of eCLIP-based RNA maps ((Van Nostrand et al. 2020)).

### Activity maps define regulatory targets

Inferring the activity map of an SF leads to a natural definition of its direct targets as exons with motifs of appropriate strength and location for regulation. As described above, the change in regulation due to knockdown for each exon is determined by the exon’s “splicing impact”, computed by multiplying predicted changes in binding by the SF’s activity map (Fig. 1f-h, Supplementary Table 2). A positive impact represents a loss of repression and increases the model’s prediction that the exon will be upregulated, while a negative impact favors a prediction of downregulation. We consider an exon to be a direct repression or enhancement target if the splicing impact of the exon is sufficiently positive or negative to improve predictions over a reduced model that excludes all splicing activity (Fig. 3a).

Comparing these predictions to the literature recovers previously described targets. For example, PTBP1 is known to silence exons in its paralogs *PTBP2* (Boutz et al. 2007) and *PTBP3* (Spellman, Llorian, and Smith 2007) during development, as well as in *RTN4* (Cheung et al. 2009) and *FLNA*(X. Zhang et al. 2016), all of which were identified as direct targets of PTBP1 by KATMAP. To assess our target predictions more comprehensively, we leveraged HNRNPC’s known role as a silencer of exons derived from the highly abundant transposable element *Alu*. These *Alu*-derived exons provide a clear ground truth for validating our target predictions. Roughly 30% of all exons upregulated upon HNRNPC knockdown are *Alu*-derived. KATMAP’s target predictions recover HNRNPC’s known silencing of *Alu*-exons, strongly sorting *Alu*-derived exons into the set of targets, with few being labelled as nontargets (Fig. 3b). That only ∼ 5% of the predicted nontargets are *Alu*-derived underscores the effectiveness of KATMAP at winnowing out these indirect effects from the SF’s true targets.

**Figure 2:**
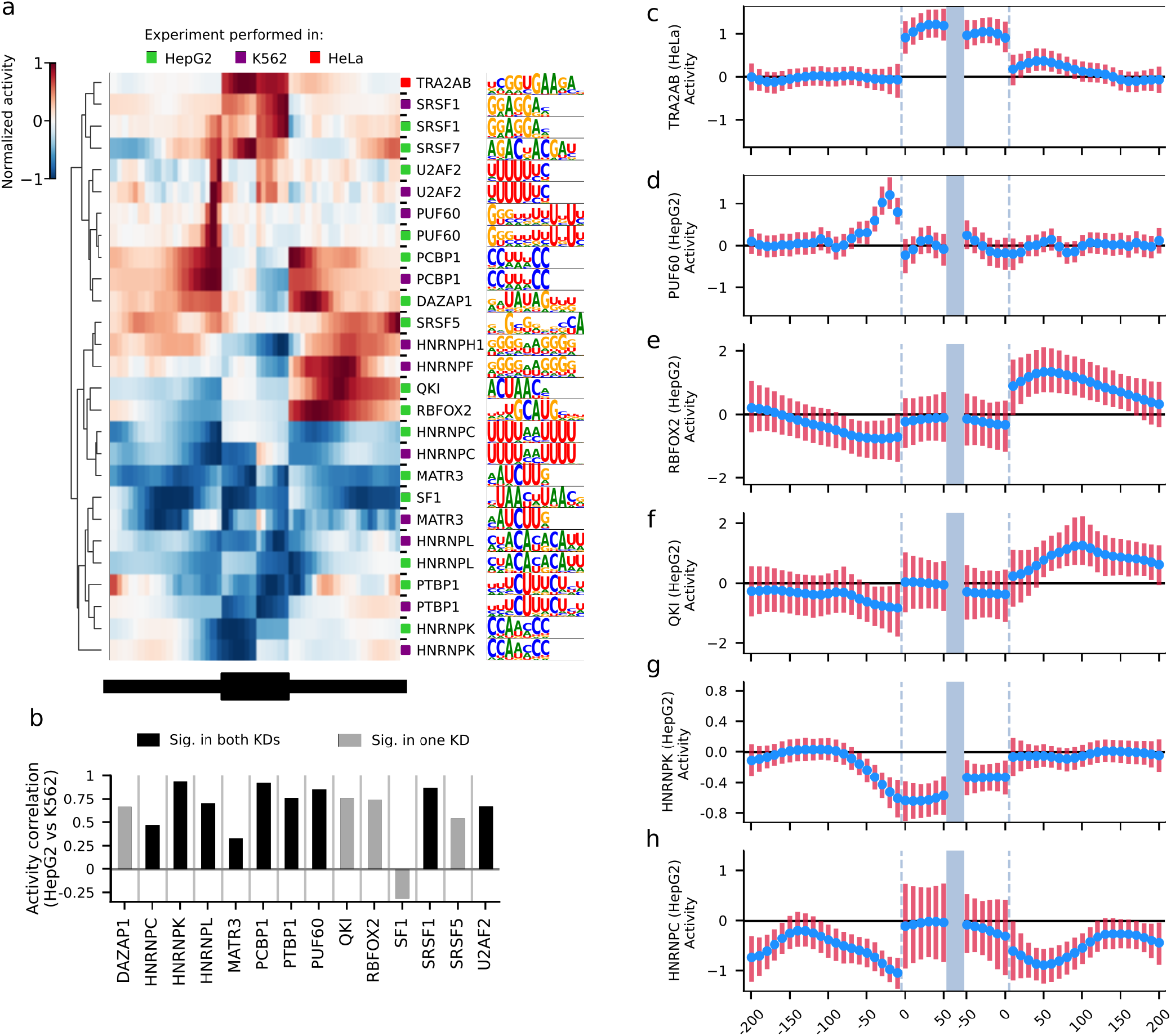
KATMAP robustly infers splicing activity. **a)** Significant activity maps and their motifs organized by hierarchical clustering. For ease of comparison, activity is normalized relative to the maximum absolute activity in each map. If the binding model included multiple motifs, only the top motif is shown.**b)** The correlation between activity maps for the same RBP inferred from different knockdowns. **c-h**) Activity maps for TRA2A/B (**c**), PUF60 (**d**), RBFOX2 (**e**), QKI (**f**), HNRNPK (**g**), HNRNPC (**h**). The blue dots indicate the posterior mean and and the lines covering the 94%-credible intervals.

### Predicted targets are bound *in vivo* and generalize across biological contexts

Knockdown and crosslinking experiments are specific to the cell types and conditions in which the experiment was performed, but if a model successfully captures regulatory activity, its predictions should generalize to other biological contexts. While our predictions are based on sequence similarity to motifs identified *in vitro* and do not incorporate crosslinking data, intersecting predicted targets with eCLIP peaks can be used to confirm binding in living cells. For the 11 SFs with credible activity maps which also had ENCODE eCLIP data, we asked whether differentially spliced target exons were enriched for *in vivo* binding compared to differentially spliced nontargets. For example, QKI is predicted to suppress inclusion upstream of the exon and enhance inclusion downstream: ∼70% and ∼80% of differentially spliced suppression and enhancing targets, respectively, had upstream and downstream QKI eCLIP peaks, compared to *<* 5% and *<* 15% of up- and downregulated nontargets (Supplementary fig. 3a). Of the 11 considered SFs, 9 showed clear enrichment for *in vivo* binding at targets, with most showing between 2-to 8-fold enrichment (Fig. 3c).

To assess whether our models can generalize across contexts, we considered SFs with knockdowns in distinct cell types and compared their predictions. For 12 of the 14 factors, models inferred from HepG2 and K562 cells yielded highly similar predictions of regulation (Fig. 3d), indicating that KATMAP consistently identifies the same underlying relationship between binding motifs and splicing regulation. To assess whether this generalizability extends across species, we applied KATMAP to mouse *Rbfox1/2/3* triple knockout data (Jacko et al. 2018). The inferred activity map (Supplementary fig. 3b) was highly consistent with those obtained from human *RBFOX2* knockdowns (Fig. 2e, Supplementary fig.3c). Using the human RBFOX models to score exons in the mouse transcriptome predicts essentially the same regulation as the splicing impacts learned directly from mouse neuronal *Rbfox1/2/3* triple-knockout data (Jacko et al. 2018) (Fig. 3e, Supplementary fig. 3d). Together, these analyses indicate that the models KATMAP infers can be applied to different tissues and torelated species, including those where *in vivo* binding data is lacking and where genes and exons not expressed in the original knockdown experiment are present.

### KATMAP identifies regulatory targets missed by differential splicing analyses

Among KATMAP’s predicted targets are exons that were not called significantly affected in the differential splicing analysis but have binding motifs properly situated to affect splicing.

**Figure 3:**
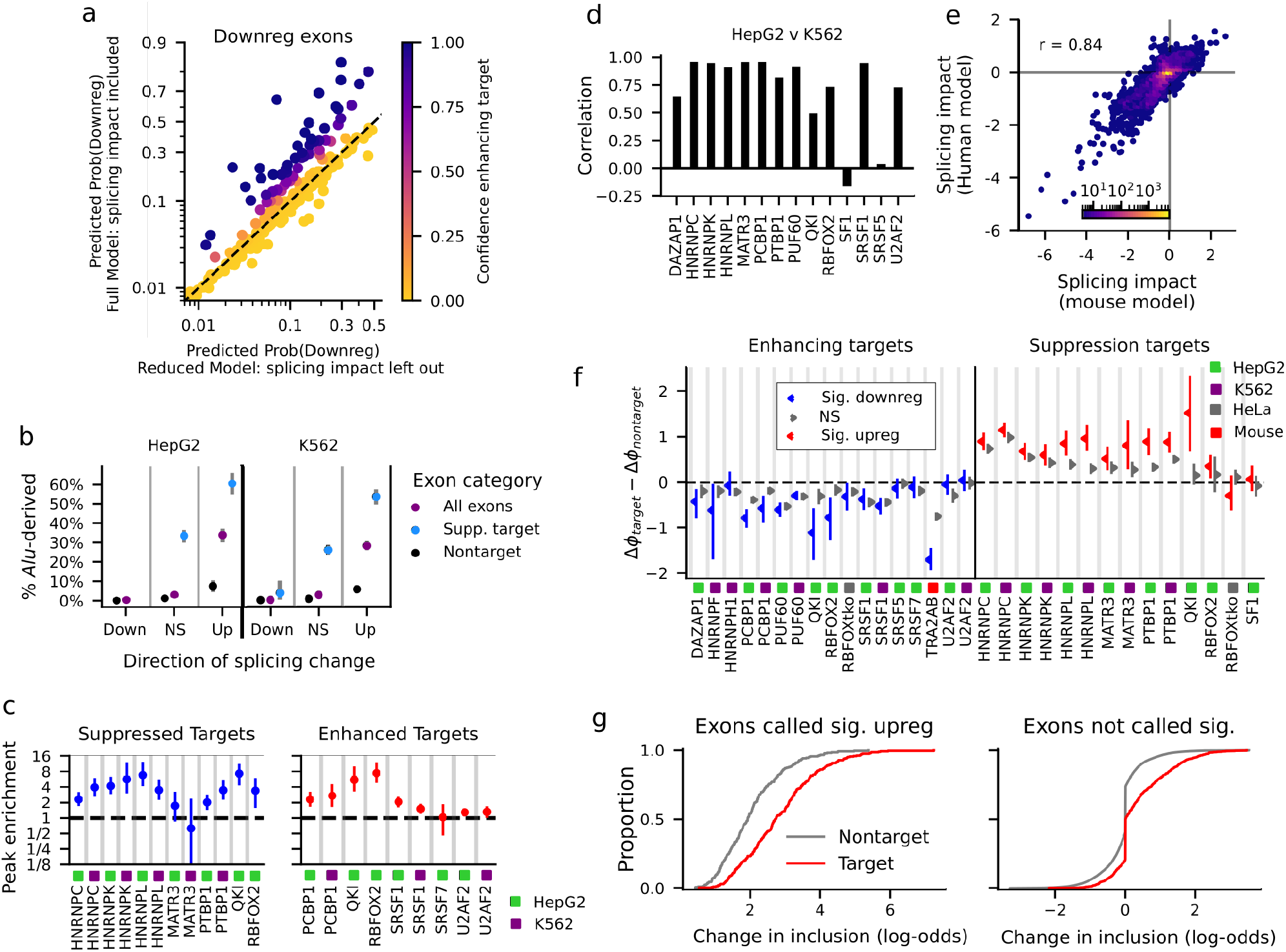
KATMAP predicts regulatory targets. **a)** The probabilities of downregulation predicted by the full model and a reduced model lacking splicing impacts for exons downregulated upon RBFOX2 knockdown (HepG2). Exons where the full model assigns a higher probability than the reduced model are likely enhancing targets, while those assigned the same predictions by both models are likely nontargets. The axes are in log-odds scale and depicted the range is 0.01–0.99. **b)** The fraction of exons derived from *Alu* among targets of HNRNPC’s suppression, nontargets, or all exons that changed in a given direction, computed for each of the three splicing categories. **c)** The enrichment of eCLIP peaks in differentially spliced targets compared to nontargets differentially spliced in the same direction. The enrichment summarizes eCLIP peaks that span the significant regions of a given SF’s activity map. **d)** The correlation in splicing impacts of exons across the human transcriptome computed from the HepG2 and K562 models for SFs with data for both cell types. **e)** Comparing RBFOX splicing impacts in the mouse transcriptome computed using the human RBFOX2 (HepG2) model and a model inferred from mouse RBFOX1/2/3 triple knockout data. **f)** The difference in average inclusion-level change between confident targets and confident nontargets among exons that were significantly altered by knockdown (red, blue) and those that were not (grey). The 94%-credible intervals were obtained via Bayesian bootstrapping. **g)** ECDF plots of the distributions of inclusion-level changes at targets and nontargets upon HNRNPC knockdown (HepG2) for exons significantly upregulated upon knockdown (left) and nonsignificant exons (right). It is the difference in the means of the target and nontarget distributions that is shown in **f**). We excluded exons where the data supported a confidently small effect of knockdown (spike at 0, |Δ*ψ*| *<* 10^−3^) to focus on those exons called nonsignificant potentially due to a lack of statistical power; the estimates with all exons included are shown in Supplementary fig. 3g

**Supplementary Figure 3:**
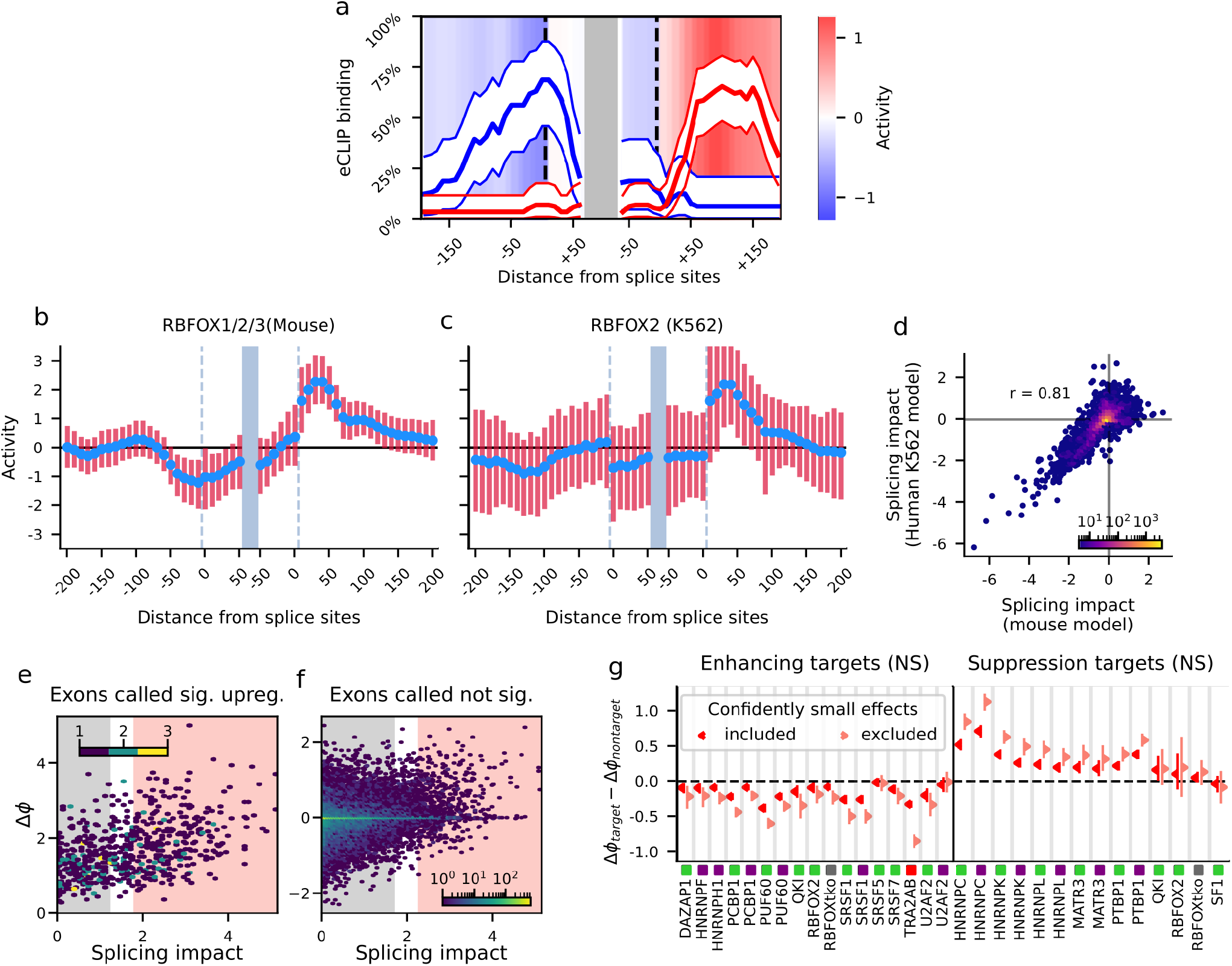
Additional validation of target predictions. **a)** The fraction of significant exons in the QKI (HepG2) analysis with QKI eCLIP peaks spanning a given position for upregulated suppression targets (blue line) and downregulated enhancing targets (red line). The regulatory activity associated with each position is indicated by the background color. **b)** The RBFOX activity map inferred from mouse *Rbfox1/2/3* knockdown data **c)** The nonsignificant human RBFOX2 activity map inferred from the K562 knockdown. **d)** Splicing impacts computed for the mouse exons using the nonsignificant human RBFOX2 (K562) model compared to those learned directly directly mouse RBFOX1/2/3 triple knockdown data. **e**,**f)** The relationship between the change in the log_2_-odds of inclusion, Δ*ϕ*, and splicing impact for exons called significantly upregulated (**e**) and those not called significantly affected (**f**) by HNRNPC knockdown in HepG2. Red and grey shaded regions indicate confident targets and nontargets respectively. **g)** The difference in average inclusion-level change between confident targets and confident nontargets among exons called nonsignificant with the spike at 0 in **f**, fig3g; |Δ*ψ*| *<* 10^−3^–included in (red) or excluded from the averages (pink). The 94%-credible intervals were obtained via Bayesian bootstrapping.

Some such exons might be true targets, for which RNAseq coverage was inadequate to confidently detect a splicing change. We asked, therefore, whether those exons predicted to be targets, but not called differentially spliced, had inclusion changes in the expected direction.

Our models were trained to predict the directions–but not the magnitudes–of changes in exon inclusion, so we first assessed whether our inferred splicing impacts were also predictive of the magnitudes of changes. We found that, among differentially spliced exons, those predicted to be direct targets had larger changes in inclusion following knockdown than nontargets (Fig. 3f). For example, among exons upregulated upon HNRNPC knockdown, those with larger splicing impacts tended to have larger changes in inclusion (Fig. 3g, Supplementary fig. 3e), with the average effect of knockdown being two-fold higher at upregulated HNRNPC targets compared to upregulated nontargets.

We find this same relationship among the exons not called differentially spliced–the exons KATMAP predicted to be direct targets changed, on average, in the expected direction compared to nontargets–consistent with the presence of true regulatory targets missed by the initial differential splicing analysis (Fig. 3f,g, Supplementary fig. 3f). An exon might not be called significantly affected by knockdown for two reasons: 1) the data provided clear evidence that inclusion did not change; or 2) the data did not provide sufficient evidence to distinguish an inclusion change from noise. The first scenario can be readily identified on the basis of confidently small ΔΨ value (Fig. 3g, Supplementary fig. 3f). Among the remaining exons, KATMAP’s target predictions are able to hone in on the subset most likely to be truly regulated by the SF, as evidenced by the propensity for change in the expected direction. By bringing to bear information about the SF’s activity and binding sites, KATMAP’s regulatory models increase the power to flag potential regulatory targets that would otherwise be ignored using conventional approaches.

### KATMAP identifies *cis*-regulatory elements and the consequences of their disruption

Implicit in KATMAP’s predictions are statements about cis-regulatory elements: for any predicted target, we can examine the collection of predicted SREs. For example, QKI is predicted to enhance the inclusion of *NF2* exon 16, which is one of two stop-codon-containing exons in the gene, the skipping of which suppresses tumor growth 3-to 4-fold in schwannoma (Sherman et al. 1997). Examining the position-wise splicing impacts suggests that QKI enhances its inclusion through a single ACUAA motif 53 nt downstream of the 5’ splice site (Fig. 4a). Other targets are predicted to be controlled by multiple binding sites. For example, HNRNPC is predicted to suppress the inclusion of a highly conserved exon in its own 5’ UTR through multiple poly-U motifs in both the up- and downstream introns (Fig. 4b).

We used this interpretability to validate selected predictions using a minigene splicing assay. We chose target exons for five splicing factors–QKI, RBFOX2, DAZAP1, PCBP1, and HNRNPC–and evaluated the effects of mutating their predicted SREs on splicing. For targets with both up- and downstream SREs, we considered the effects of mutating only the upstream or only the downstream elements. We inserted the reference and mutated exons into reporters and measured exon inclusion (Fig. 4c). To confirm that the effects of mutation were mediated by the expected SF, we assayed reporters with and without knockdown of the expected factor.

**Figure 4:**
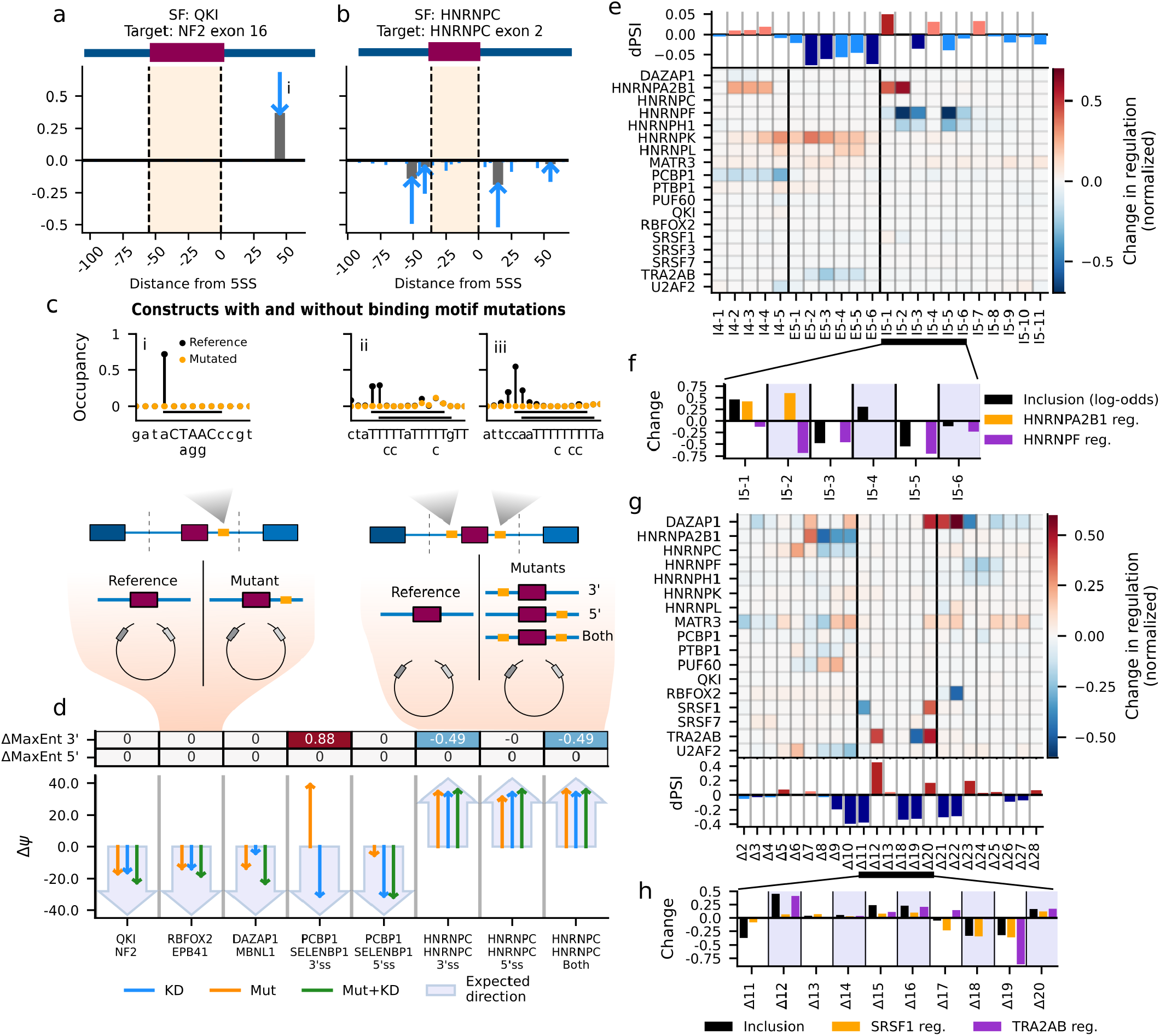
KATMAP predicts the consequences of disrupting cis-elements. **a**,**b)** The expected changes in regulation due to loss of QKI (**a**) and HNRNPC (**b**) binding in predict targets. The blue arrows indicate the change in regulation from the pre-knockdown levels (tail of arrow) to the post-knockdown (grey bar). **c)** Schematics of the minigene reporter design. **Top:** The expected occupancy at the predicted splice regulatory elements (i–iii from **a**) and the predicted effects of mutating these elements. The reference sequence is on the X-axes with the expected binding motif capitalized; the specific mutations are below. **Bottom:** The reference and mutated versions of predicted targets were inserted into a splicing reporter. Inclusion was assayed with and without SF knockdown. **d)** The results of the reporter assay. The large arrow depicts the direction the mutations were expected to alter inclusion. The thin arrows depict the observed effect of knockdown, mutation, and mutation+knockdown. **e)** The reported changes in HRAS exon 5 inclusion (top) after ASO treatments and the changes in SF regulation predicted by KATMAP (bottom). Changes in regulation are normalized to the highest activity position in each SF’s activity map. Black vertical lines separate the exon from the introns. **f)** Zoomed in view of altered inclusion and regulation by HNRNPF and HNRNPA2B1 in the downstream intron. **g)** The reported effects of deletions on inclusion of the TRA2B poison exon (bottom) and KATMAP’s predicted changes in regulation (top), as in **e. h)** Zoomed in view of the effect of deletions in the exon on inclusion and predicted regulation by TRA2A/B and SRSF1, based on the 200 nt exonic models.

Inclusion changed in the expected direction for seven of the eight mutant reporters (Fig. 4d). Mutation of the QKI and RBFOX2 binding sites downstream of the *NF2* and *EPB41* exons, respectively, reduced inclusion comparable to knockdown. Similarly, disrupting the DAZAP1 binding sites upstream of the *MBNL1* exon decreased inclusion as expected, but to a greater extent than the knockdown. The PCBP1 downstream mutations decreased inclusion as expected, but to a smaller degree than the knockdown, possibly because the three upstream PCBP1 sites remained intact. The upstream PCBP1 mutations are the one case where the inclusion change was opposite to our expectation. In this case, the mutations occurred very close to the 3SS and are predicted to increase the strength of the PPT, which may override any impact of the loss of PCBP1 binding. In the HNRNPC reporters, both the upstream and downstream HNRNPC mutations behaved as expected, with either being sufficient to increase inclusion about 40% to near complete inclusion. Thus, with the exception of one of the PCBP1 mutants, splicing changes always occurred in the expected direction.

To further evaluate KATMAP’s cis-element predictions we asked whether our models could be used to interpret two previously published tiling assays: an antisense oligonucleotide (ASO) tiling of *HRAS* exon 5 (X. Chen et al. 2023) and a deletion tiling of *TRA2B* ‘s poison exon (Leclair et al. 2020). The *HRAS* exon tiling experiment found that ASOs targeting the downstream intron led to decreases in exon inclusion by blocking enhancing activity of HNRNPF/H proteins at *G*_3_ and *G*_4_ motifs (X. Chen et al. 2023) as well increases in inclusion consistent with a previously reported silencing element just downstream of the 5SS, likely mediated by HNRNPA1 (Guil et al. 2003). While the HNRNPF and H1 knockdowns both yielded significant KATMAP activity maps, the HNRNPA maps failed to pass our model comparison criteria. However, the *HNRNPA2B1* (K562) knockdown yielded an activity map supporting intronic repression (Supplementary fig 4a) and *HRAS* exon 5 was the model’s highest confidence repression target, so we included it in these analyses.

For each SF, we predicted the changes in regulation due to ASOs blocking SREs, assuming the ASOs disrupt recognition of their complementary nucleotides and one additional base on either side. The SFs predicted by KATMAP to be most strongly affected by the ASOs were HNRNPF and HNRNPA2B1, with HNRNPH1 showing a profile similar to HNRNPF (Fig. 4e). Two of the three ASOs predicted to disrupt HNRNPF’s enhancing activity led to decreased inclusion and 4 of the 5 ASOs predicted to disrupt HNRNPA2B1’s repression led to increased inclusion (Fig. 4e,f). The exception was ASO I5-2, which was predicted by KATMAP to disrupt both HNRNPA2B1’s repression and HNRNPF’s enhancement and yielded no change in inclusion (Fig. 4f), perhaps because blocking these activities canceled each other out. The success of the HNRNPA2B1 model in explaining the effects of ASOs both suggests that it captures real regulatory activity and that our model comparison criterion might be excessively stringent in some cases.

Analysis of a deletion tiling experiment of the *TRA2B* poison exon again implicated expected factors. KATMAP primarily explains the effects of exonic deletions on inclusion in terms of the SR proteins TRA2A/B and SRSF1 (Fig. 4g). While deletions that increase inclusion are typically interpreted as removing silencing elements, KATMAP instead predicts that these deletions promote positive regulation by TRA2A/B by moving its binding sites closer to the splice sites, where activity is higher. The *TRA2B* poison exon is particularly long (276 nt), while our activity maps only extend 60 nt from the splice sites. To confirm that this window size does not bias our predictions we refit SRSF1 and TRA2A/B models using larger windows that extend up to 200 nt into the exon (Supplementary fig 4b,c). These extended models infer that TRA2A/B and SRSF1 activity decays with distance from the splice sites, as previously demonstrated for SR proteins(Graveley, Hertel, and Maniatis 1998), and similarly predict that the deletions increase regulation by moving binding sites nearer the splice sites (Fig. 4h, Supplementary fig 4d-g). Finally, we performed a permutation analysis– exchanging ΔΨ among deletions within the exon, and within the upstream, and downstream introns. The observed associations for TRA2A/B and SRSF1 were consistently stronger than the chance associations under permutation (*p* = 0.003 and *p* = 0.007, respectively).

**Supplementary Figure 4:**
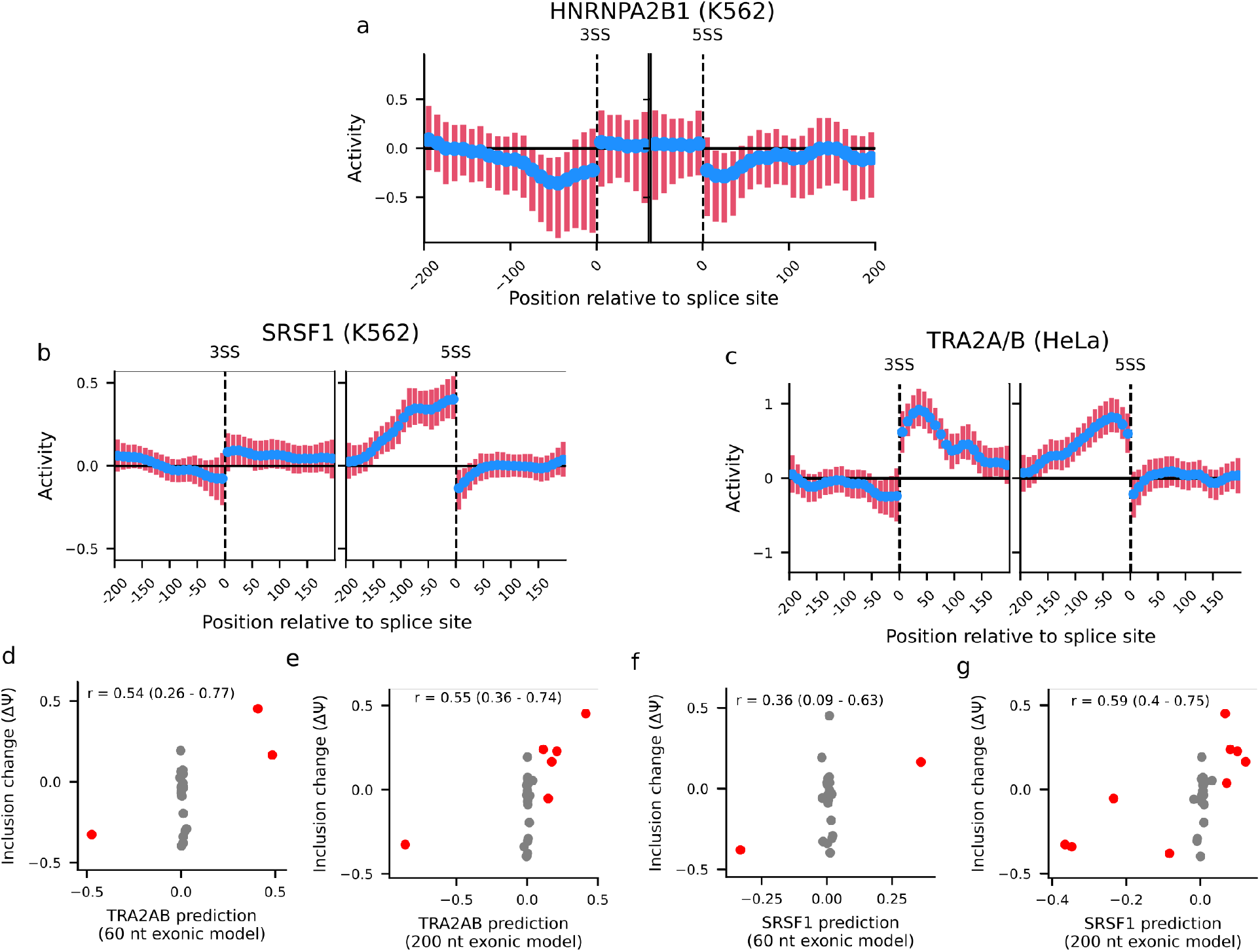
**a-c)** The activity map from the HNRNPA2B1 (K562) ENCODE knockdown (**a**) using the standard 60 nt exonic window, and activity maps inferred from **b)** SRSF1(K562) and **c)** TRA2A/B knockdowns, estimating an expanded activity for positions 200 nt up- and downstream of splice sites. **d-g)** Association between change in exon inclusion (Leclair et al. 2020) and KATMAP predicted changes in TRA2A/B and SRSF1 regulation based on the standard activity maps (**d, f**) and extended 200 nt activity maps (**e, g**). Deletions predicted to change regulation by more than *±*0.05 are colored red. Credible intervals for the correlations were computed using Bayesian bootstrapping.

Together these observations support both KATMAP’s ability to infer cis-regulatory elements responsible for splicing regulation and to predict the effects of their disruption.

### Target predictions reveal cooperative regulation

Our model makes the simplifying assumption that SFs act independently of each other. However, we reasoned that we could uncover evidence of cooperative regulation by examining our model’s predictions for deviations from this null assumption. To test this possibility, we focused on RBFOX and QKI, which have strikingly similar activity maps despite binding distinct motifs (Fig. 2e,f), raising the possibility of a functional relationship between them. Earlier analyses noted that the sets of exons responsive to QKI and RBFOX knockdown are overlapping (Brosseau et al. 2014; Li et al. 2018; Van Nostrand et al. 2020), and observe-denrichment of QKI eCLIP signal at RBFOX-downregulated exons. However, clear evidence of QKI and RBFOX binding at the same exons was not observed, and it was concluded that the correlated effects of knockdown did not reflect direct coregulation by the factors (Van Nostrand et al. 2020), though another study found binding of both factors at a subset of exons (Li et al. 2018). These changes could arise without direct coregulation, for example if perturbing RBFOX impacted QKI activity, altering splicing of its targets. These analyses, however, could not distinguish between the direct and indirect effects of knockdown, a limitation that might have obscured signatures of coregulation.

We used KATMAP’s ability to discriminate direct regulatory targets from indirect effects to reassess the case for direct coregulation. In the scenario of strong cooperativity, knocking down either factor should primarily affect exons bound by both factors, whereas exons bound by only one should be minimally affected. As KATMAP models one SF at a time, an exon with a binding motif for the knocked down factor will be called a target, whether or not a motif for the coregulator is present. Cooperative regulation should produce a clear signature at predicted targets: knockdown-affected targets of each factor should be enriched for targeting by both coregulators compared to targets unaffected by knockdown (Fig. 5a,b). We detected this expected signature of cooperativity, with differentially spliced QKI targets being enriched for RBFOX enhancing regulation (Fig. 5c). The reciprocal relationship was also observed: RBFOX2 targets affected by *RBFOX2* knockdown are enriched for QKI binding sites (Fig. 5d). Further, this pattern is evident in mouse cells depleted of all three RBFOX paralogs (*Rbfox1/2/3*) (Fig. 5e), supporting that coregulation of exons by RBFOX and QKI proteins is conserved in mice (∼90 mya divergence).

**Figure 5:**
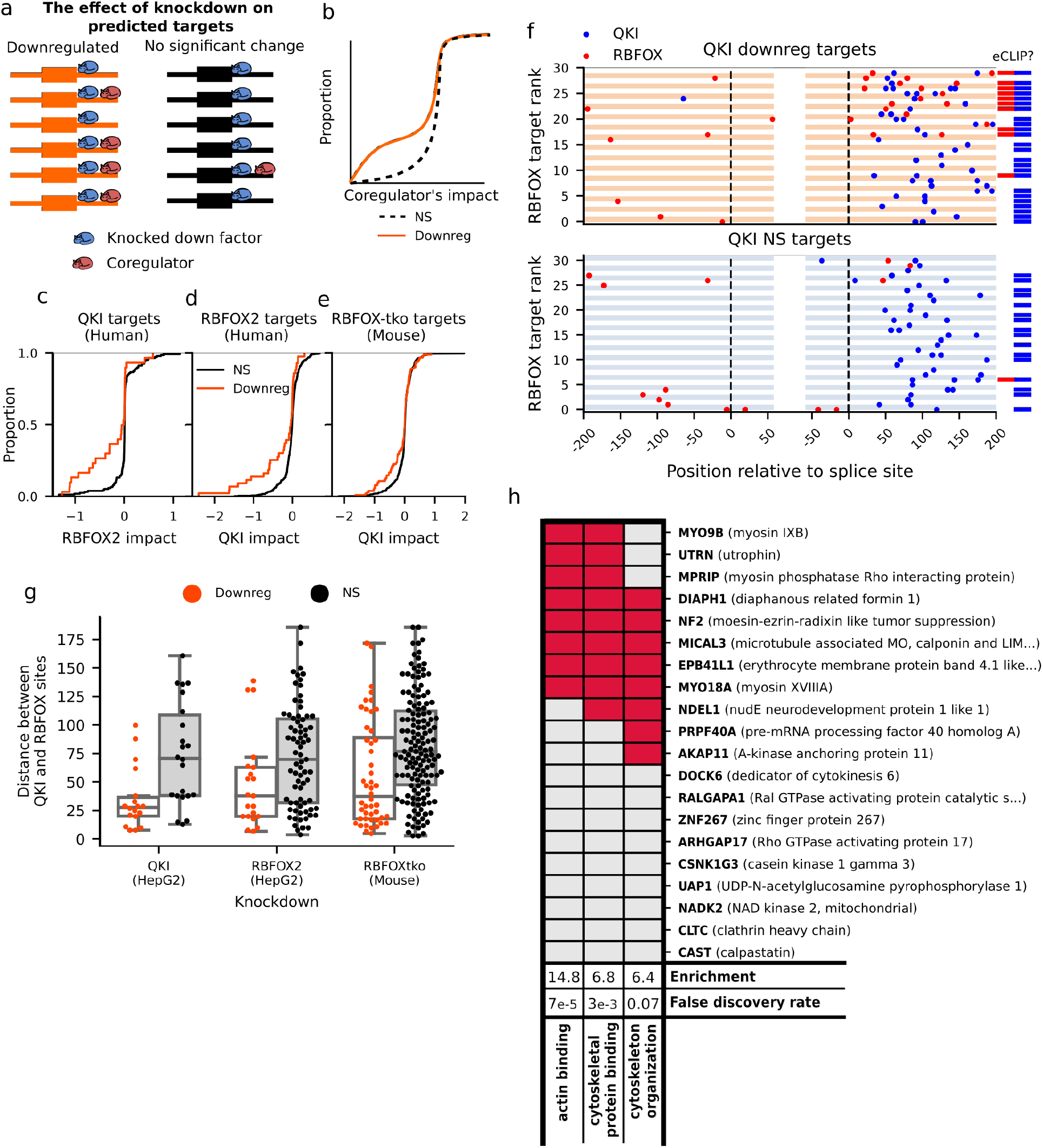
QKI and RBFOX cooperatively regulate a splicing subprogram. **a)** In the case of cooperativity, predicted targets affected knockdown should be enriched for the coregulator compared to those unaffected by knockdown. **b)** The expected distributions of the coregulator’s splicing impact computed at KD-affected (orange) and KD-unaffected targets (black). **c-e)** Distributions of RBFOX2 (**c**) and QKI (**d**,**e**) splicing impacts at the targets of the other protein (**c**: QKI; **d**,**e**: RBFOX), comparing the KD-affected (orange) to KD-unaffected targets (black). **f)** Locations of predicted binding sites (|rel. change-in-binding| *>* 20%) for both RBFOX (red) and QKI (blue) in KD-affected QKI enhancing targets affected (**top**) and an equal number of KD-unaffected targets (**bottom**). The exons are sorted by RBFOX splicing impact and the bars to the right denote the presence of eCLIP peaks in the downstream intron. **g)** The distributions of distances between the nearest QKI and RBFOX sites in KD-affected (orange) and unaffected (black) targets. **h)** GO-analysis of confident QKI-RBFOX cotargets; red denotes GO-membership.

These coregulated exons have both QKI and RBFOX binding sites (Fig. 5f)–frequently multiple of each–occurring as clusters of interspersed motifs separated by a few dozen nt (Fig. 5g). This proximity of binding sites suggests that QKI and RBFOX proteins could physically interact, which is supported by prior yeast two-hybrid (Lang et al. 2021) and pull-down mass spectrometry data (Huttlin et al. 2021). Further, reassessment of the eCLIP data focused on direct targets supports binding of both QKI and RBFOX to the same exons: QKI crosslinking is detected at both knockdown-affected and unaffected QKI targets, but RFBOX2 is primarily bound at the targets affected by QKI knockdown. Together, these observations support direct, cooperative coregulation of exons by QKI and RBFOX proteins.

Our analyses identified twenty human exons with binding sites for both QKI and RBFOX proteins and responsiveness to either *QKI* or *RBFOX2* knockdown (Supplementary Table 3). Of these twenty, eleven are in genes involved in cytoskeleton binding and organization, with seven being actin-binding proteins (Fig. 5h). This represents not only strong enrichment for these categories relative to expression-matched control genes, and relative to the full set of RBFOX or QKI targets, consistent with a focused subprogram active only in cells where both QKI and RBFOX are expressed and distinct from their individual regulatory roles. Cytoskeletal remodelling and cell migration are integral to early neuronal development, and a number of these exons occur in genes with known roles in neural and glial biology. For example, *NDEL1* is critical for cell migration during neural differentiation (Youn et al. 2009) and *NF2* exon 16 skipping strongly promotes growth of glial tumors (Sherman et al. 1997). Both RBFOX and QKI proteins are expressed in neuronal progenitor cells (NPCs), which differentiate into both neurons – where RBFOX predominates – and glia – where QKI is predominant – suggesting a potential venue for coregulation (Hardy 1998; X. Zhang et al. 2016).

### KATMAP’s models uncover the SFs responsible for splicing changes in RNAseq data

The set of splicing changes observed in RNAseq datasets following perturbations such as stress, drug treatments, or cell differentiation are expected to result from changes in the activities of splicing factors. Identifying these causal factors would help to elucidate the underlying regulatory network and prioritize targets for intervention. While the ENCODE RBP knockdowns offer a resource to explore this, the splicing changes in the knockdowns represent mixtures of direct and indirect effects. Indeed, most knockdowns caused abundant splicing changes in both directions, even for established activators (e.g. SRSF1) and repressors (e.g. HNRNPL) of splicing, where direct targets should be down- and upregulated respectively. We infer that most splicing changes in the ENCODE knockdowns result from indirect effects, with less than 30% of splicing changes directly caused by loss of the knocked down RBP (Fig. 6a). The high proportion of indirect effects likely results from the multi-day period of antibiotic selection common to knockdown experiments, and the extensive cross regulation among splicing factors.

KATMAP distills knockdown/RNAseq data into predictions about direct regulation across the transcriptome. If a given SF contributed to the splicing changes in an RNAseq dataset, the splicing impacts learned from analyses of its knockdown data should be predictive of which exons were affected. We express this logic with a multinomial logistic regression that predicts each exon’s splicing changes based on the splicing impacts inferred from the different ENCODE knockdowns (Fig. 6b). To validate this approach, we used the splicing models inferred from K562 knockdowns to explain splicing changes in the HepG2 knock-downs, and *vice versa*. This approach correctly identified the depleted SF as most predictive in 21 out of 23 knockdowns (91%) (Fig. 6c), demonstrating that our regulatory models can generalize beyond their training data to infer the SFs underlying transcriptomic changes.

In most knockdown experiments, this analysis additionally implicated secondary SFs as responsible for some indirect effects (Fig. 6d,e, (Supplementary Table 4)). For example, in the *HNRNPC* K562 knockdown, while splicing changes were most consistent with a loss of HNRNPC activity, they also implied a loss of PCBP1 and DAZAP1 regulation (Fig. 6d). This loss was likely driven by decreased levels of *PCBP1* and *DAZAP1*, whose mRNAs were significantly downregulated in the *HNRNPC* knockdown. More generally, *PCBP1* was significantly downregulated (*FDR <* 0.2) in 42 knockdowns and upregulated in 5. This variation in *PCBP1* levels drives indirect effects across the ENCODE knockdowns: the knockdowns where *PCBP1* was secondarily downregulated had splicing changes consistent with loss of PCBP1 regulation, and *vice versa* in cases where it was upregulated (Supplementary fig. 5a), with a significant Pearson’s correlation of 0.3 (94%-CI: 0.2–0.4). The presence of measurement error causes Pearson’s *r* to underestimate the underlying association (Supplementary fig. 5a,b). Employing an error-aware hierarchical model that disentangles the true association from the estimation errors revealed a larger correlation between *PCBP1* expression and splicing variation of 0.55 (94%-CI: 0.41–0.69)(Supplementary fig. 5b, c),.

These analyses provide a proof-of-principle that splicing factor perturbations can be inferred using KATMAP’s models in the context of well-controlled experiments. To assess the potential of this approach in more complex and disease-relevant contexts, we applied KATMAP to a published comparison of splicing changes between primary and metastatic pancreatic ductal adenocarcinoma (PDA) samples, which uncovered a role for RBFOX proteins in repressing metastasis (Jbara et al. 2023). Unlike knockdown data, cancer samples typically represent a mixture of distinct etiologies, which may obscure the signal of any specific causal factor. Indeed, the original analysis did not recover unambiguous RBFOX motifs, but rather amalgams of G- and C-rich motifs, some of which had similarity to RBFOX secondary motifs. Applying KATMAP, we directly inferred loss of RBFOX regulation in the metastatic samples through a single analysis (Fig. 6f), without the need to first identify motifs and then match them to putative SFs. We further infer a gain of SRSF1 regulation, a factor known to promote tumorigenesis, (Karni et al. 2007) which was missed in the original analysis. The inferred perturbations to RBFOX and SRSF1 are consistent with reported protein level changes in these samples, with RBFOX protein levels decreased in the metastatic lineages and SRSF1 protein levels elevated (Jbara et al. 2023).

## Discussion

Since the development of RNA sequencing, perturbation experiments have been the predominant tool for probing the rules by which regulatory factors control gene expression and processing. But these studies have typically reported visual summaries of regulation, which are not directly useful for quantitatively interpreting regulation in new biological settings. Further, most analyses do not distinguish between the direct and indirect effects of perturbation, instead assuming that the signal from direct targets will rise above the background of indirect effects. Our approach, KATMAP, extracts actionable insights about splicing regulation from SF perturbation data. The activity maps KATMAP generates are not only interpretable visualizations of an SF’s regulatory activity but also models that can be applied to make predictions in new contexts. These predictions distinguish the direct from indirect effects of SF perturbation, allowing indirect effects to be further interpreted in terms of secondary factors and preventing them from contaminating conclusions drawn about the perturbed factor’s biological role. Our approach is supported by the reproducibility of inferred activities and predictions when applied across cellular contexts and related species. The model design makes KATMAP a general framework for rigorously applying insights from perturbation data to new types of analyses.

**Figure 6:**
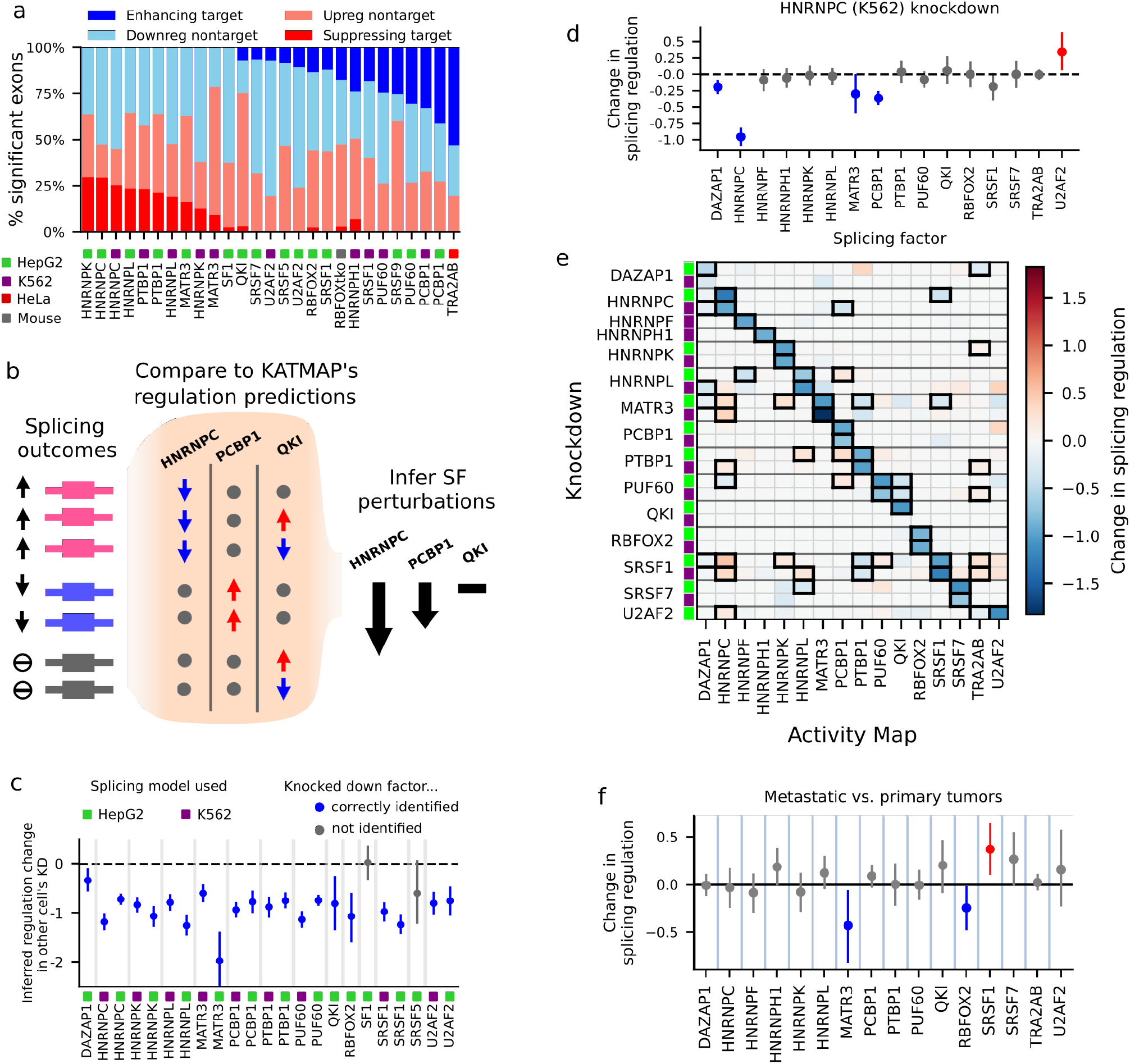
KATMAP’s regulatory models uncover SF perturbations. **a)** The fractions targets and nontargets among differentially spliced exons (rMATS FDR *<* 0.05). Ambiguous target predictions (*Pr*(*Target*) and *Pr*(*Nontarget*) *<* 0.8) are not included. **b)** Schematic showing how we infer underlying SF perturbations by comparing splicing changes to KATMAP’s regulatory models. In this example, exons predicted to be suppressed by HNRNPC are upregulated and exons enhanced by PCBP1 are downregulated, suggesting that decreases in HNRNPC and PCBP1 regulation caused the splicing changes. There is no association between the splicing changes and QKI’s predicted targets, suggest no perturbations to QKI’s regulation. **c)** Predicted changes in regulation for the knocked down factors, inferred by comparing to all activity models learned from the other cell type. Knockdowns where the expected factor gave the strongest signal are indicated in blue. **d)** The predicted regulation changes in the HNRNPC (K562) knockdown, with 94% credible intervals. **e)** Predicted regulation changes as in **d** for knockdowns of all RBPs with a significant activity map, using ASHR to enforce sparsity. Significant changes in regulation are denoted with thick boxes. **f)** Predicted differences in SF regulation between metastatic and primary PDA samples.

**Supplementary Figure 5:**
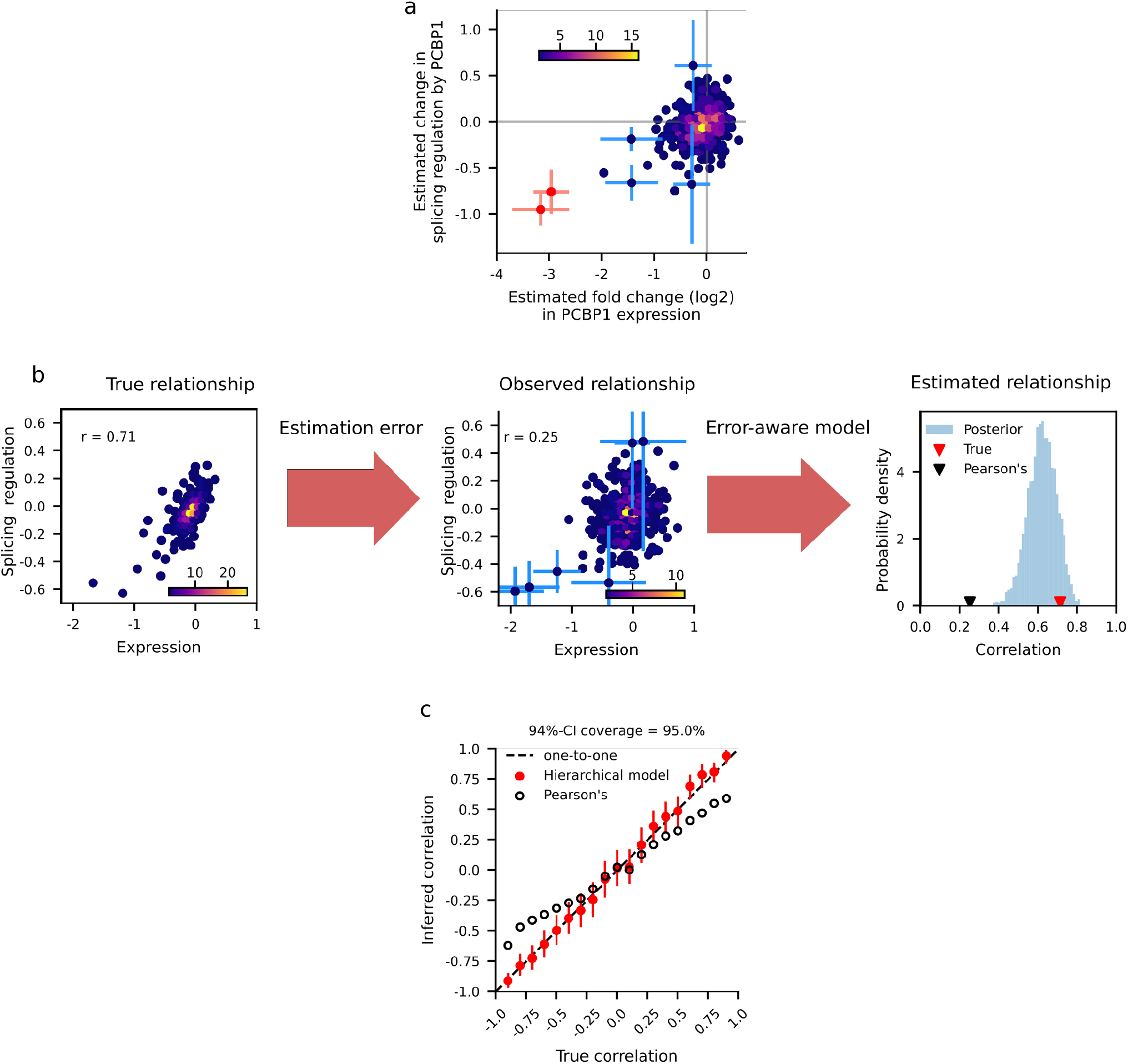
Deconvoluting true associations from measurement error. **a)** Comparison across ENCODE RNAseq datasets between inferred perturbations to PCBP1’s splicing regulation and PCBP1’s expression variation. The PCBP1 knockdowns are in red. The error bars show the uncertainty in the point estimates (94%-ci) for select datasets. **b)** A true association (left) is obscured by measurement error (middle). Pearson’s correlation computed from the noisy point estimates substantially underestimates the true association. If the degree of uncertainty associated with each point estimate (blue lines) is known, it can be incorporated into a hierarchical model to recover the true association (right). **c)** Evaluation of the hierarchical applied to simulated data with true correlations between -0.95 and 0.95 and the same measurement errors as the PCBP1 analysis. The hierarchical model recovers the true association, with a nearly one-to-one relationship between the estimates and true values. Pearson’s correlation applied directly to the point-estimates, however, underestimates the association by ∼ 50%.

Despite these advantages, our approach has several limitations. First, we make the simplifying assumption that each SF can be modelled individually and ignore potential co-operative regulation between SFs. Nonetheless, the regulatory models learned by KATMAP can be employed as null hypotheses to uncover evidence of cooperativity, as seen for RBFOX and QKI proteins. Second, because we designed KATMAP to predict regulation from sequence, we can only learn regulatory models for factors with known sequence specificity. This currently limits inference to SFs with available *in vitro* binding data, but we are actively exploring alternatives such as learning binding models from crosslinking data. Finally, while based on a general model of regulation, the model is currently limited to describing splicing regulation. Recent work suggests that KATMAP’s assumption of position-specific activity also applies to transcription factors (TFs) (Duttke et al. 2024), suggesting that modified versions of KATMAP might find application in modeling transcription.

In distilling knockdown experiments into generalizable models, KATMAP provides the opportunity to leverage decades of experimental work to answer new questions. The insights our model extracts from perturbation data will benefit work focused on specific regulatory factors, providing information about activity and targets, and quickly generating hypotheses to further probe that factor’s biology. More generally, many researchers who generate RNAseq dataset will perform a differential splicing analysis, typically uncovering hundreds of splicing differences, which are challenging to interpret. Integrating KATMAP into RNAseq workflows provides a means to focus downstream analyses on particular factors responsible for the phenotype of interest. Our reanalyses of pancreatic cancer data illustrates a use case for inference of SF drivers of splicing changes in a clinical dataset (fig6f). Identifying the SFs and SREs underlying splicing programs in disease may speed the search for therapeutic targets. Further, because KATMAP’s predictions are explicitly framed in terms of changes in binding at *cis*-elements, our models can facilitate interpretation of sequence variants. This is highlighted in our minigene experiments where we successfully identified regulatory sequence elements and designed mutations that disrupt that regulation. Similar analyses have applications to ASO design by providing information on which *cis*-elements are likely to change inclusion in the desired direction. Focusing on these regions can reduce the need for laborious tiling or deletion assays during therapeutic development, and inference of specific SFs may aid in design of ASOs active in relevant tissues.

## Online Methods

### Selecting the knockdown experiments

We selected ENCODE knockdown RNAseq experiments for proteins annotated as splicing factors (Van Nostrand et al. 2020) and which had a good affinity model derived from RBNS (Lang et al. 2021) using RBPamp (Jens et al. 2022) or a PWM derived from RNAcompete (Ray, Kazan, Chan, et al. 2009) obtained from the CISBP-RNA data database (Ray, Kazan, Cook, et al. 2013). We use the rMATS differential splicing (Shen et al. 2014) results for the hg19 human reference genome in our analyses.

### The structure of the KATMAP regression model

The goal of KATMAP is infer a SF’s regulatory activity and targets from knockdown data by explaining the resulting splicing changes in terms of changes in SF binding. Accomplishing this requires several inputs. First, we need the results of a differential splicing analysis, which assigns labels *Y* to exons indicating whether their inclusion increased, decreased, or was unaffected by knockdown. These are the observations KATMAP seeks to predict. The second input is a model of the SF’s sequence specificity. This assigns exons in the differential splicing analysis a set of scores, *S*, which evaluate all potential binding sites within predefined windows around their splice sites. These scores, *S*, are what KATMAP will use to explain the splicing changes, *Y*, in terms of SF binding, and ultimately to infer regulatory activity. Finally, additional information about each exon can be included using a set of optional linear predictors, *X*. In our analyses, we use these linear features to control for expression and pre-knockdown exon inclusion levels.

### Defining the splicing changes, *Y*

Any differential splicing analysis that labels each exon as unchanged, up-, or downregulated may be used to define the observations, *Y*. We use one-hot encoding to define *Y*_*i*_ as (1, 0, 0), (0, 1, 0), or (0, 0, 1) if exon *i* is labelled downregulated, unchanged, or upregulated, respectively.

In our analyses, we defined splicing changes by further processing the rMATS outputs with ashr to better account for the uncertainty in rMATS’s estimates (Stephens 2017). This step had two motivations. First, ashr accomplishes something like a multiple test correction on parameter estimates, but accounting for the possibility that more exons are upregulated than downregulated (or vice versa). Second, while KATMAP does not consider the magnitude of splicing changes, some of our downstream analyses do. The shrinkage estimates provided by ashr control for measurement error due to low read counts and yield a cleaner view of splicing differences than the raw rMATS Δ*ψ* summaries.

Briefly, we used the read counts and rMATS p-values to reconstruct the effect size in log-odds scale, which we denote Δ*ϕ*, and associated uncertainty (Extended Methods). We then pass these values to ashr to obtain shrinkage estimates for Δ*ϕ* and associated standard deviations. We construct 94% credible intervals based on these, calling exons differentially spliced if the 94%-CI does not overlap zero and nonsignificant otherwise.

### Defining the training set

We use the rMATS skipped exon table in our analyses. If a set of skipped exons partially overlapped or shared flanking exons, we retained only the most significant to avoid double counting. We include all remaining significant exons in our training sets. For computational efficiency, we downsample the nonsignificant exons, retaining 2,000 randomly chosen exons.

### Scoring motifs with a binding model

To explain splicing in terms of binding, KATMAP requires some model of the perturbed SF’s binding motif. The binding model may be a traditional position-weight matrix (PWM) obtained from RNAcompete, one or more position-specific affinity matrices (PSAM) (e.g. from RBPamp), or more general affinity models. The key requirement is that the binding model approximately orders potential binding sites from weakest to strongest. In our analyses we used PSAMs derived from RBNS (Lambert et al. 2014) using RBPamp (Jens et al. 2022) when available, and PWMs derived from RNAcompete (Ray, Kazan, Chan, et al. 2009) obtained from the CISBP-RNA database (Ray, Kazan, Cook, et al. 2013). We excluded RBNS models where the PSAMs showed strong complementarity to the sequencing adapters, which likely reflects a technical artefact rather than true sequence specificity.

To score a *k*-length motif *x* with a single 4-by-*k* PWM or PSAM, *W*, we sum the scores (in log-scale) for each nucleotide in the motif

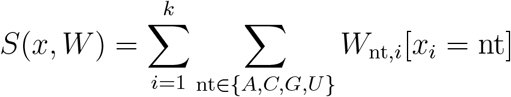

where the Iverson bracket [*x* = nt] equals 1 if true and 0 if false.

If we have multiple matrices *W*_1_, .., *W*_*n*_, each with associated weights *a*_1_, .., *a*_*n*_, then we compute the combined score as

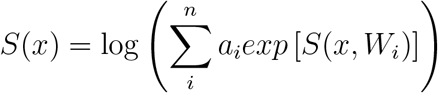

In practice, we always define the scores such that the optimal motif is assigned a score of 0 and all other motifs receive negative scores. This is accomplished by subtracting the score of the optimal motif from all raw scores.

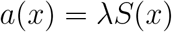

Even for affinity models, we do not assume that these initial sequence-based scores are fully accurate representations of affinity. Rather, we treat them as related to log-affinity, but potentially differing by some scaling factor. To account for this, we include an *affinity-scale* parameter, *λ*, which rescales the binding scores.

These rescaled scores are used as log-affinities from which occupancy and changes-in-occupancy are computed. If the scores are truly linearly related to log-affinity, this can be thought of as calibrating the scores against the observed splicing changes. If scores are related to affinity in a monotonic but not linear manner, this rescaling can be thought of widening or narrowing the interval of scores assigned appreciable changes in occupancy.

### Scoring the sequences around exons

KATMAP scores sequence in windows around the splice sites of each exon. In our analyses, we scored 200 potential binding sites on the intronic and 60 on the exonic side of the splice sites of each cassette exon. If desired, larger or smaller windows can be used, or the regions adjacent to the flanking exons’ splice sites can be included.

Our model calculates a log-affinity for each of the potential binding sites analysed per exon. For computational tractability, rather than model binding at nucleotide resolution at each exon, we aggregate neighboring binding sites into non-overlapping regions of ten potential binding sites each. We index these regions, 1..*J*, where in most of our analyses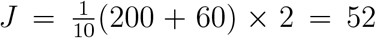. More generally, in an analysis with a different region size, different intronic or exonic window size, or number of the splice sites (e.g. including the flanking exons)

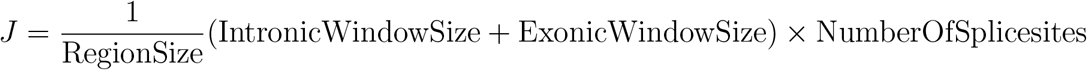

We then assign each subregion *j* of exon *i* a log-affinity. We denote the motif scores of the ten binding sites ending in subregion *j* as **S**_*ij*_ and compute the log-affinity of the region by first scaling the log-affinity of each binding site by *λ* and taking the log of the sum of their exponents:

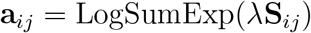

### Defining the binding function

For a given free protein concentration *F*, the occupancy of a site with given log-affinity, *a*, is described by the Langmuir equation

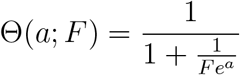

The change in occupancy, then, is determined by the free protein concentration before and after knockdown/overexpression (Supplementary fig. 6a). If the free protein concentration changes by some factor *k*, the change in occupancy can be computed (Supplementary fig. 6b)

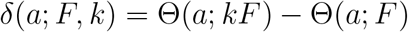

However, these parameters, *F* and *k*, while biophysically meaningful, are not immediately interpretable in terms of which affinities experience large changes in binding. We therefore reparameterize the change-in-binding function in terms of its shape. First, we include a most-relevant-affinity parameter, *m*, indicating which affinity experienced the greatest change in binding. Second, we describe the magnitude of the change in binding at sites with affinity *m* with a *greatest-change-in-binding* parameter, *d* (Supplementary fig. 6b,c). That is, we reparameterize such that that maximum change in occupancy, *d*, occurs at log-affinity *m*.

The corresponding biophysical parameters can be computed by solving

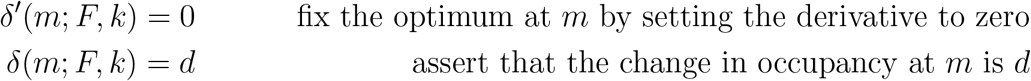

which yields

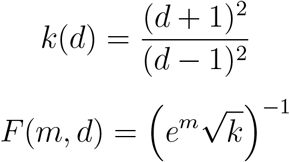

so

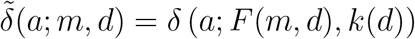

where 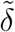 denotes a reparameterization of *δ*.

Our model’s predictions are constructed by multiplying these binding changes by other parameters we term *activity coefficients, α*. Because *d* changes the height of the binding function but not the relative shape (unless *d* is near 1 or −1,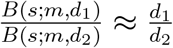; Supplementary fig. 6c), we cannot usually identify both the magnitude of binding changes and the magnitude of regulatory activity: most effects on the model’s predictions of changing *d* from value *d*_1_ to *d*_2_ can be entirely reversed by scaling the magnitudes of the activity coefficients by 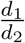. To resolve this, we normalize the changes in binding by |*d*| to compute the relative change-in-binding (Supplementary fig. 6d)

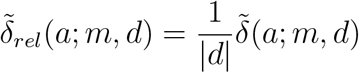

Further, because for all values of |*d*| *<* 0.95 yield essentially the same relative changes-in-binding, we fix 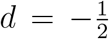 for knockdown experiments and 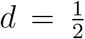 for overexpression. If |*d*| *>* 0.95, as might be the case for knockout data, the relative change-in-binding function flattens out, but this can be approximated with small values of the affinity scale parameter, *λ* ≪ 1 (Supplementary fig. 6e)

To recap, we score the sequences around each exon’s splice site using a model of the RBP’s binding site. We then divide the sequence around each exon into non-overlapping 10 nt regions, each containing ten potential binding sites. We denote the set of all scores as *S*, where *S*_*ij*_ refers to the scores of the ten potential binding sites in region *j* of exon *i*. We then scale these scores with an *affinity scale* parameter, *λ*, and treat the resulting values as log-affinities. We then compute the log-affinity of the regions based on the affinities of the ten binding sites contained in each. Finally, we assign each region a relative changing-in-binding, determined by the *most-relevant-affinity* parameter, *m*. We describe this entire procedure of transforming the motif score to relative changes-in-binding with the binding function

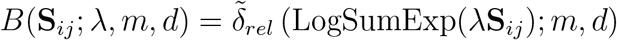

fixing 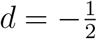 for knockdown experiments.

**Supplementary Figure 6:**
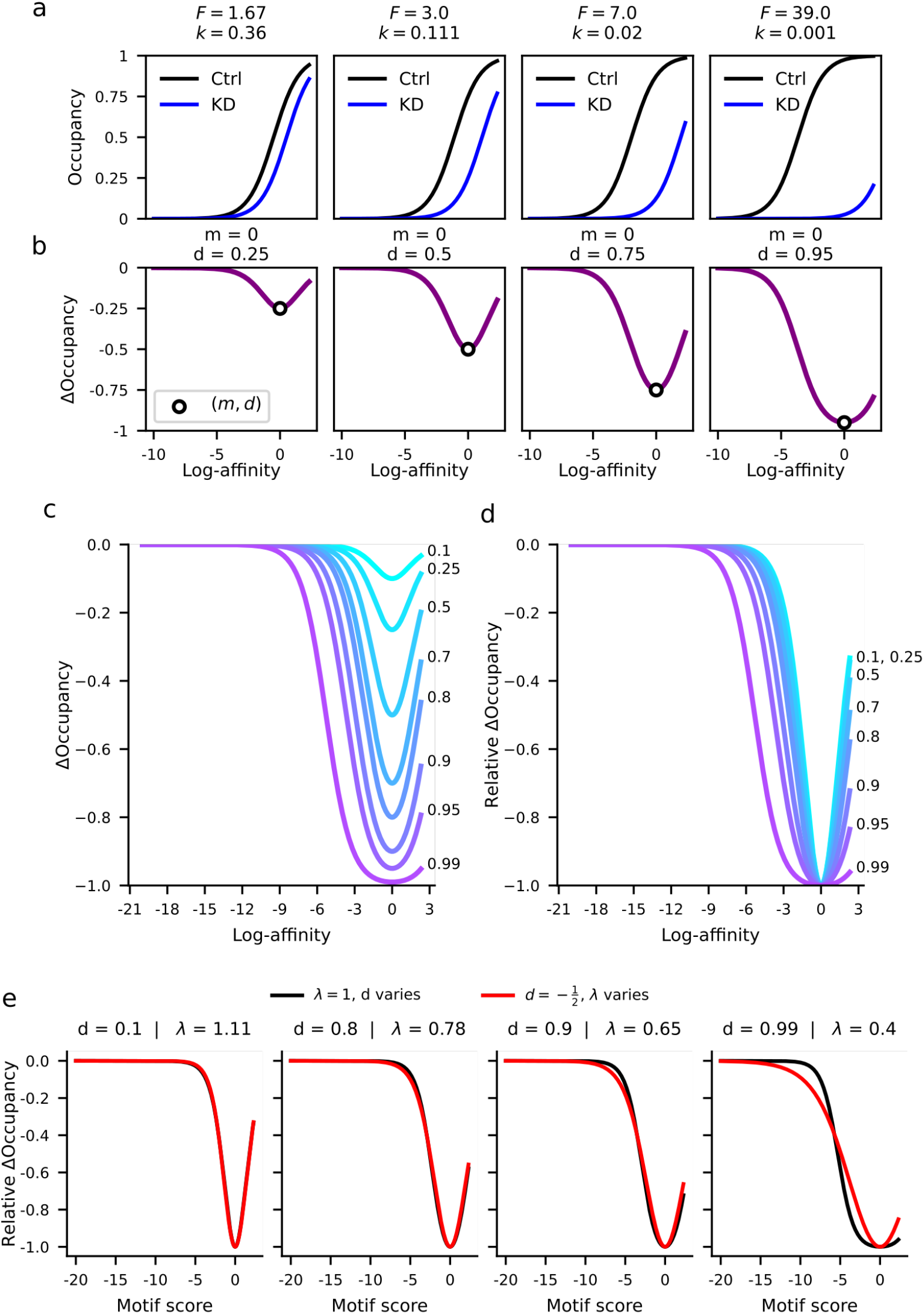
Binding functions. **a)** The affinity-occupancy relationship before (black) and after knockdown (blue) for different pre-knockdown free protein concentrations (*F*) and knockdown efficiencies (1 − *k*). **b)** The change in binding functions that result from occupancy curves in **a)**, with the corresponding values for the shape parameters *m* and *d*. Note that unlike the biophysical parameters (*F* and *k*) these shape parameters explicitly define the affinity and magnitude of the greatest binding change (open circle). **c-d)** Absolute change in binding, 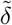, (**c**) and relative change in binding, 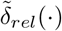, (**b**) with the most-relevant affinity *m* = 0 for different values of *d*. **e)** The best approximations of the binding function at a given *d* obtainable by rescaling the affinities

### Predicting changes in regulation from changes in binding

We describe the position-specific effect of RBP binding with a set of *activity coefficients*: *α*_1..*J*_. A positive value of *α*_*j*_ indicates that binding at region *j* enhances exon inclusion. A negative value instead describes suppressing activity. Critically, we assume that the activity varies smoothly with position, described later when discussing our model’s priors.

To compute the predicted effect of loss of binding near exon *i* on its splicing outcome, we multiply the change in binding at each region *j, B*(*S*_*ij*_; *λ, m, d*), by the corresponding activity coefficient *α*_*j*_. We make the simplifying assumption that activity is additive across all binding sites, so the total change in activity due to the changes in binding across regions 1..*J* is

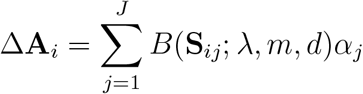

### Connecting changes in regulation to observed splicing changes

To predict the observed splicing changes, the model must assign each exon *i* three probabilities, the probability of being downregulated, unchanged, or upregulated: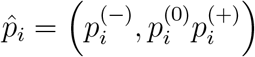.

We describe the log-odds 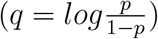 of these probabilities with an additive model:

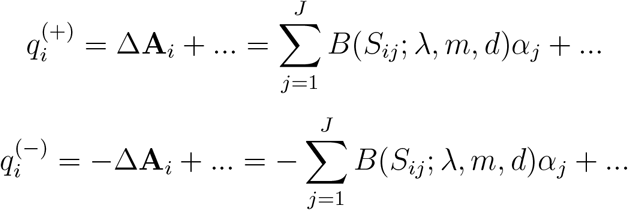

That is, if the activity at exon *i* increases, the predicted log-odds of inclusion being upregulated increases and the log-odds of being downregulated decreases. In an overexpression experiment, the changes-in-binding *B*(*S*_*ij*_; *λ, m, d*) are positive, and upregulation is driven by gains in enhancing activity and downregulation by gains in suppressing activity. In knock-down experiments, the changes-in-binding are negative, flipping the signs of the change in activity: upregulation is driven by loss of suppressing activity and downregulation by loss of enhancing activity.

### Incorporating additional information into the predictions as linear features

Our models aims to explain the results of the differential splicing analysis in terms of SF binding, but other information unrelated to binding is likely to be predictive of which exons were called differentially spliced, for biological and/or technical reasons. We incorporate these features by adding a standard linear model into the predictions. The *K* linear features included in the model are organized as an *I*-by-*K* table, *X*. All features are mean-centered and standardized. Any quadratic terms are created using the standardized features to reduce the dependence structure of the target posterior distribution.

Because not every linear feature will be relevant to each of the three predictions–downregulated, unchanged, upregulated–we divide the linear features *X* into three (potentially overlapping) subsets, denoted *X*^(−)^, *X*^(0)^, and *X*^(+)^. For example, the read count of an exon influences the statistical power to detect changes in splicing and so should inform the predicted probability of being unchanged, *p*^(0)^ and so is assigned to *X*^(0)^. The original inclusion level of an exon determines whether up- or downregulation is detectable, because there is little room for inclusion to increase or decrease when the exon was already nearly 100% or 0% included. So, the inclusion-level prior to SF perturbation is likely directly informative of 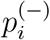 and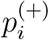, and so is assigned to both *X*^(−)^ and *X*^(+)^. These feature is incorporated into the predictions as

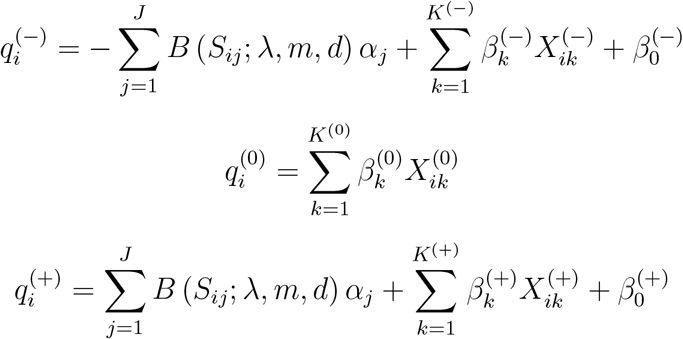

where 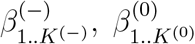, and 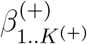 denote the linear coefficients describing the effect the linear features on the predictions, and where *β*^(−)^ and *β*^(+)^ are the intercepts.

### Connecting predictions to observed splicing changes

The probabilities for category *z* are computed as

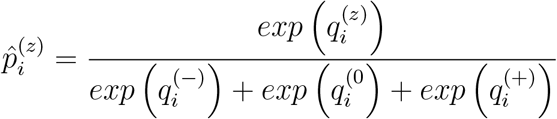

We need to connect these to the observed splicing changes with a categorical distribution

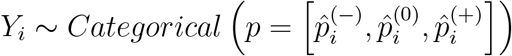

The probability of an observation under this distribution is simply equal to the predicted probability for that category. For example if exon *i* is downregulated, the likelihood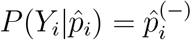.

### Defining the model’s priors

We represent assumptions about our model’s parameters with Bayesian priors. Some of the priors we employ reflect what we consider as plausible based on our understanding of molecular biology. Others reflect statistical considerations that aim to avoid unrealistically strong predictions or complications to statistical inference. For example, given the limitations of experimental data and modelling thereof we do not believe KATMAP should ever make extremely confident predictions on the order of million-to-one odds. In other cases, we wish to avoid identifiability issues, where many combinations of distinct parameter values would yield essentially the same predictions. As we describe the model’s prior, we will note for each parameter which consideration is at play.

### Priors on the binding parameters: *λ, m*

The parameter *λ* scales the motif scores *S* to assign binding sites log-affinities *S* × *λ*. When *λ <* 1 the scores are compressed, assigning a narrower range of affinities. When *λ >* 1 the scores are stretched, assigning a wider range of affinities. We choose a prior that asserts both to be equally plausible. Beyond this, the prior we place on *λ* largely reflects a practical consideration about operations in log-scale. Multiplication on log-scale corresponds to exponentiating the actual affinities:

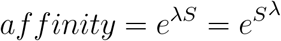

Even modest values of *λ* will yield extremely large or small affinities. So we constrain *λ* to a relatively narrow range of values by using a unit-normal prior

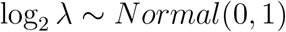

which places about 99% of the prior probability on *λ* between 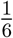 and 6 (Supplementary fig. 7a).

The most-relevant-affinity parameter *m* describes the which affinity experiences the greatest change in binding. Assuming the binding site model is accurate, motifs with moderate to high affinity should experience changes in binding; motifs with very low affinity are too weak to be appreciably bound in the first place. So a biologically reasonable prior would favor values of *m* closer to the affinity of the optimal motif and disfavor very low affinities. But beyond this, it is difficult to precisely express how plausible a given value of *m* is.

However, *m* determines how much SF occupancy our model predicts was lost from near each exon. It is easier to assess the biological plausibility of these statements. Certain values of *m* might imply that the average exon has *>* 20 copies of the SF bound within a few hundred nucleotides. For SFs that bind specific sequence motifs, this is unrealistically high. So rather than explicitly define a prior on *m*, we instead define our prior on the average change in binding near each exon

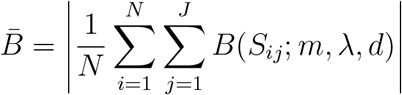

We divide this quantity by the number of potential binding sites per exon (*J* = 52 in our analyses) and place the prior on the log-odds of this fraction :

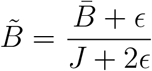

(small constants, *ϵ* = 10^−7^, are added to the numerator and denominator to avoid numerical issues due to computational precision). We then define our prior as

**Supplementary Figure 7:**
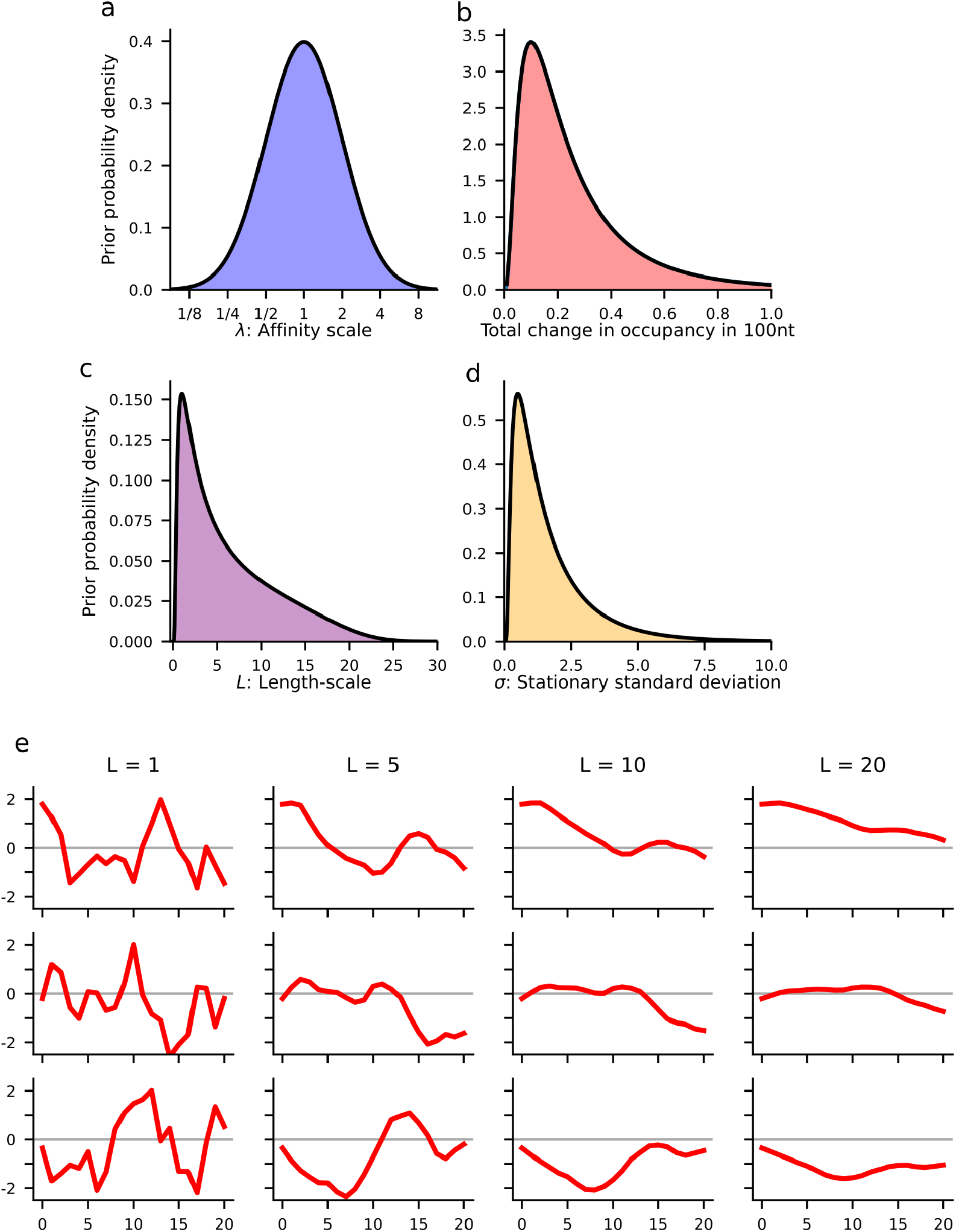
Priors on the hyperparameters and resulting spatial correlations. Prior densities on **a)** the affinity scale (in *log*_2_-scale); **b)** the average amount of binding lost per exon (implicit prior on most-relevant-affinity); **c)** the length-scale of the Gaussian process prior (in units of 10 nt regions) on the activity coefficients; and **d)** stationary standard deviation of the Gaussian process. **e)** Examples of the spatial smoothness resulting from different length-scales.

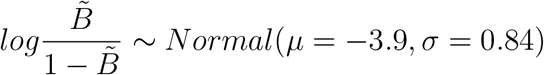

which places 99% of the prior probability on the average change in binding per exon between 0.12 and 7.8 (Supplementary fig. 7b). To exclude extremely unrealistic values of *m*, we constrain the model to only consider values between the median log-affinity and the maximum affinity per 10nt region *log*(10) (or more generally *log*(*RegionSize*))

### Assuming that activity varies smoothly with distance: *α*_1..*J*_

A fundamental assumption of our model is that the effect of SF binding on exon-inclusion depends on where the protein is bound relative to the exon. This is described by activity coefficients *α*_1..*J*_. We further assume that the activities of nearby regions, say *j* = 2 and *j* = 3, are similar, but that distant positions may have very different activities. Statistically, we express this with a Gaussian process prior that assumes the activity coefficients are spatially correlated (Rasmussen and Williams 2006). Gaussian process can be conceptualized as a multivariate normal distribution, where the covariance between variables *α*_*j*_ and *α*_*k*_ is a decreasing function of how far apart they are |*j* − *k*|.

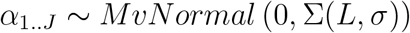

We choose the smooth, yet flexible, Matern-3/2 covariance function to express these spatial correlations (Rasmussen and Williams 2006). Because we further want to assume that the effects of exonic versus intronic binding may be very different, we restrict the spatial correlations to pairs of regions within the same biological window. We express this with a set labels *R*_1..*J*_ that indicate whether region *j* is on the intronic side of the 3’ splice site, the exonic side, the exonic side of 5’-splice site, or the intronic side. If a pair of regions *j* and *k* correspond to differ biological labels, the spatial correlation between *α*_*j*_ and *α*_*k*_ is set to zero

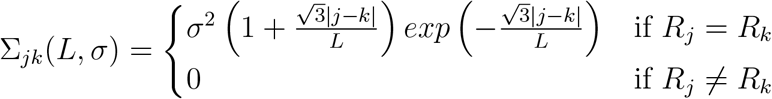

Assuming this spatial correlation structure requires introducing two additional parameters that must be learned. The length-scale, *L*, determines how far apart a pair of regions *j* and *k* must be before they are weakly correlated. The stationary standard deviation, *σ*, determines the range of magnitudes the activity coefficients are expected to span. To prevent the inference algorithm from wasting computation by exploring large but practically equivalent regions of parameter space, we constructed priors that avoid both large and small values of these hyperparameters (Supplementary fig. 7c,d, see Extended Methods). This places 99% of the prior probability for the length-scale on the interval 3.2 nt *< L <* 226 nt (Supplementary fig. 7c), which ranges from near independence to correlations that span the whole intronic window (Supplementary fig. 7e). For the stationary standard deviation, 99% of the prior probability falls in the interval 0.14 *< σ <* 8.7 (Supplementary fig. 7d).

### Assumptions about the linear coefficients: *β*_0..*K*_

Because we mean center and standardize the linear features, *β*_*k*_ indicates how much an increase in linear feature *k* by one standard deviation changes the log-odds of the prediction. Because the impact is in log-odds scale, coefficients of magnitude |*β*| *>* 10 would have extreme impacts on the prediction: increasing the linear feature by one standard deviation would increase a predicted probability of *p* = .01 to *p >* .99. As the standardized features typically span a range of more than four standard deviations, we employed a weakly-informative prior for the linear coefficients and intercept, using a normal density with mean 0 and standard deviation 5

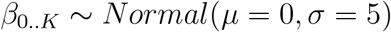

The model is not sensitive to this choice of prior, as it spans the range of reasonable effect sizes, both strong and weak.

### High-level overview of model inference

The goal of Bayesian inference is to determine which parameters are plausible given the observed data. In practice, this is often a harder task than formulating the statistical model itself. Evaluating the exact plausibility requires applying Baye’s theorem, which expresses a reasonable proposition: to determine how plausible a given hypothesis *h* is, you must compare it to every other hypothesis admitted by the model, *h* ∈ Θ:

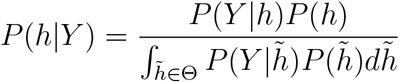

This is typically impossible to compute exactly, because it requires integrating over the entire parameter space. In the case of KATMAP’s regression model, this parameter space has *>* 50 dimensions.

A common solution is Monte Carlo sampling. Rather than evaluate the exact posterior density *P* (*θ, ϕ* |*Y*), we instead seek to obtain a sample of hundreds of parameter values which follow the posterior distribution

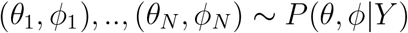

This posterior sample provides a representative set of parameters that are plausible given the observed data from which we can estimate the posterior mean. We can also construct credible intervals describing the range of plausible parameters values. Because the model predictions for each exon are computed based on these parameters, it is further possible to obtain interval estimates for all of the exon-level predictions.

### Bayesian inference with the KATMAP model

When writing the posterior for KATMAP’s model it is useful to divide the model parameters into two groups. First, we have the regression coefficients *θ* = (*α*_1..*J*_, *β*_0..*K*_). Second, with slight abuse of terminology, we have the hyperparameters *φ* = (*λ, m, L, σ*). Partitioning the parameters in this manner makes clear the hierarchical nature of the model

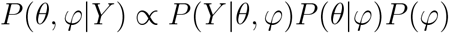

Integrated nested Laplace approximations (INLA) provide a means to leverage this structure and divide the problem of posterior inference into two manageable, nested sub-problems. First, an inner step focuses on identifying which regression coefficients are plausible given the data and hyperparameters. This approximates the conditional posterior *P* (*θ*|*Y, φ*) for a specific hyperparameter value, *φ*. Second, an outer step focuses on determining which hyperparameters are plausible given the data. This approximates the marginal posterior for the hyperparameters *P* (*φ*|*Y*)

### Approximating the conditional posterior over the regression coefficients

The inner step takes advantage of how the model simplifies if the hyperparameters are held fixed to a particular value. The known binding parameters (*λ, m*) would transform the table of motif scores *S* into a fixed table of changes-in-binding *B*_(*λ,m*)_. The known spatial correlation parameters would yield a fixed prior on the activity coefficients. The problem reduces to a generalized linear model, albeit a Bayesian formulation with priors.

We use second-order optimization identify the optimal regression coefficients as the mode of the conditional posterior, 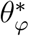, determining the step size with a backtracking line search (Nocedal and Wright 2006). This requires evaluating the gradient ∇ and Hessian matrix *H*, which we compute using automatic differentiation with the jax library (Bradbury et al. 2018). How sharply the log-posterior is curved around the mode is described by the Hessian matrix 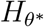 and can be used to describe the uncertainty about the coefficients with a covariance matrix, 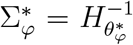. This provides a multivariate normal approximation to the conditional posterior

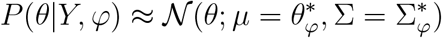

### Approximating the marginal posterior over the hyperparameters

The outer step evaluates the plausibility of specific hyperparameters *φ* by approximating the marginal posterior, *P* (*φ*| *Y*). The hyperparameters, *φ* only relate to the observations through the regression coefficients *θ*. So evaluating the plausibility of some specific *φ* requires considering all possible values the regression coefficients *θ* could take. Rue, Martino, and Chopin (2009) accomplish this using the normal approximation to the conditional posterior obtained in the inner step evaluated at its mode, 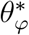

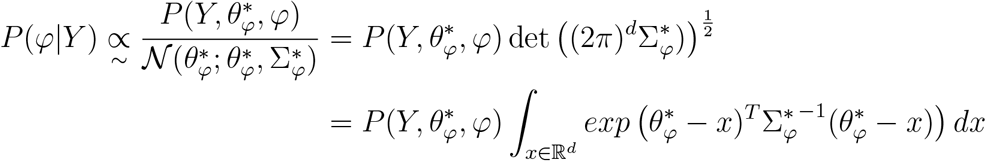

This approximation to the marginal density multiplies the joint density 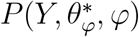 by the volume of regression coefficients that are plausible given *φ*.

### Monte Carlo sampling from the joint posterior with INLA

We couple adaptive importance sampling with these integrated nested Laplace approximations (INLA) to draw samples of plausible parameters from the joint posterior. First, we use adaptive importance sampling and the INLA approximation to draw a representative sample of plausible hyperparameters from their marginal posterior, *φ*_1_, .., *φ*_*N*_ ∼ *φ*|*Y*. This explores the hyperparameters in a way that iteratively hones in on the true posterior, by fitting mixture model approximations with importance-weighted expectation-maximization (Cappé et al. 2008) (Supplementary fig. 8 a, Extended Methods). Second, for each sampled hyperparameter we solve the resulting conditional model by optimization, approximating the conditional posteriors (including their correlation structure) over the regression coefficients with a multivariate Gaussian. We then sample regression coefficients from these conditional posteriors, *θ*_1_, .., *θ*_*N*_ ∼ *θ*|*Y, φ*_1..*N*_. Together, these constitute an approximate sample from the joint posterior, (*θ*_1_, *φ*_1_), .., (*θ*_*N*_, *φ*_*N*_) ∼ *θ, φ*|*Y*. We run this algorithm twice for each knock-down experiment, generating two separate posterior samples. If both runs converge to the same distribution, we consider inference successful and merge the distributions.

### Evaluating convergence

We use two criteria to assess whether our Monte Carlo samples are reliable representations of the target posterior. First, we use the split 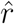-statistic (Vehtari, Gelman, et al. 2021) to assess whether our two inference runs converged to the same distribution. Values of 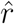 close to one indicate highly overlapping distributions and if marginal distributions of all parameters yield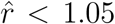, we consider the Monte Carlo sampling runs to have converged. Second, during adaptive importance sampling of the hyperparameters, we stabilize the importance weights with Pareto-smoothing. This automatically provides a diagnostic statistic, 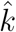, indicating whether the resulting importance sample is reliable. We use the cutoff suggested by Vehtari, Simpson, et al. (2024), requiring 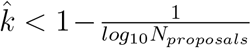 or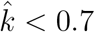 –whichever is more stringent– for the posterior samples to be considered trustworthy.

### Evaluating evidence of splice regulatory activity

To quantify evidence of in favor of splice regulatory activity, we compare to a reduced model with the splicing impacts removed. We compare the full and reduced models using Pareto-smoothed importance sampling leave-one-out cross-validation (PSIS-LOO) (Vehtari, Simpson, et al. 2024), a fully Bayesian approach to model comparison which uses the posterior samples from the full and reduced models to estimate whether the full model better predicts held out data. We bootstrap observations to compute a Z-score describing our confidence that including splicing activity in the model improves performance.

### Determining which activity maps are significant

We consider an activity map significant based on two criteria. First, we require that at least two activity coefficients be significantly nonzero (98%-CI does not overlap zero). Second, we require a model comparison z-score *>* 2.

**Supplementary Figure 8:**
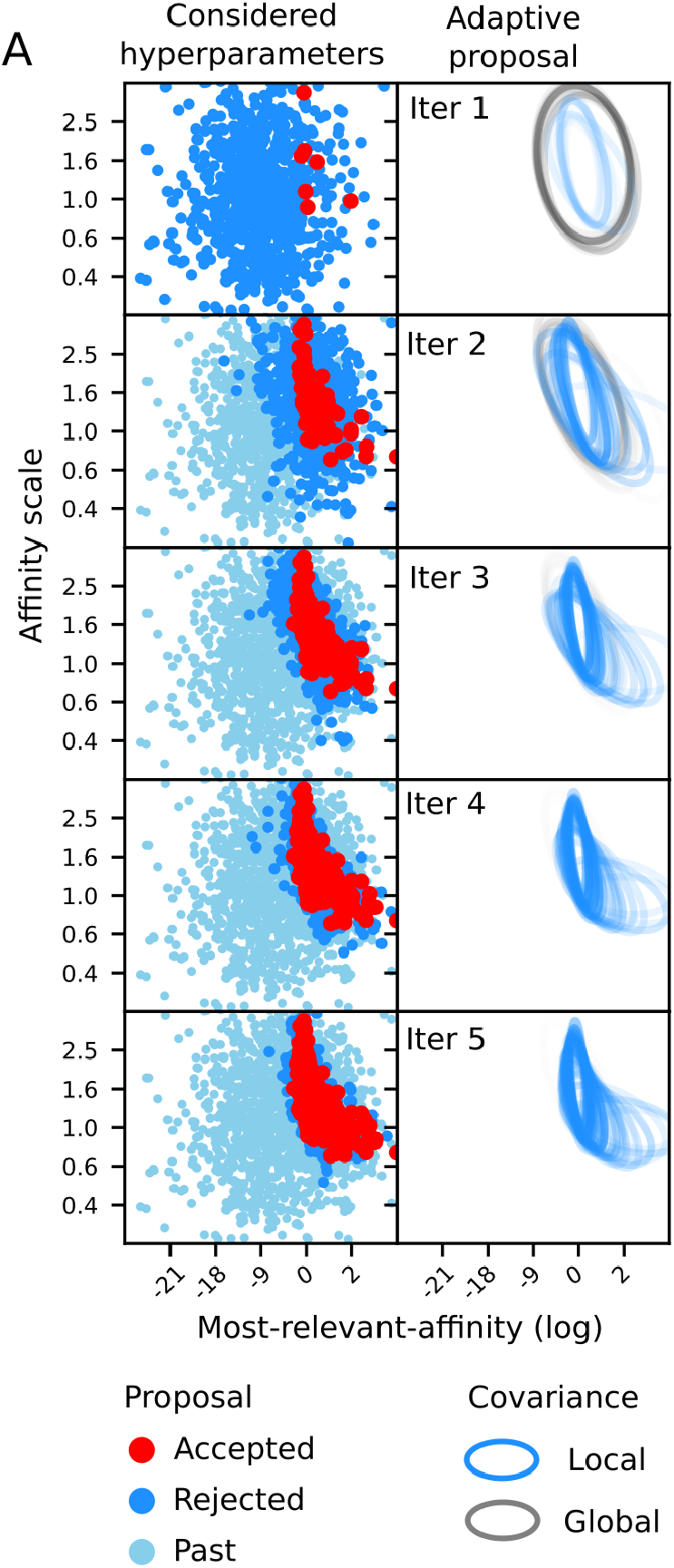
Posterior inference for the KATMAP model. **a**) Exploration of the hyperparameters through adaptive importance sampling, showing the evaluation points (left) and the adaptive proposal distribution at successive iterations. New hyperparameters (blue) are drawn from the proposal estimated in the previous iteration, which are in turn used to refine the adaptive proposal distribution to better match the target posterior. The proposal is a student T mixture model fit by importance-weighted Expectation-Maximization. Upon termination, hyperparameters are accepted by importance sample to obtain a sample from the posterior (blue).

We additionally considered whether the splicing changes are well-described by the expected motif. We excluded any activity models that choose binding parameters which assigned an unreasonable number of binding changes (*>* 20 per exon). This indicates that the knockdown-affected exons are distinguished by some characteristic sequence composition, but it is not well described by the perturbed factor’s expected motif. This excluded SRSF9 from our analysis, which yielded a significant map by assigning appreciable occupancy to most sites in the transcriptome.

### Evaluating a reduced model without splicing activity

To evaluate evidence of splice regulatory activity, we compare to a reduced version of the model that only includes the linear predictors. The predictions under this model are computed

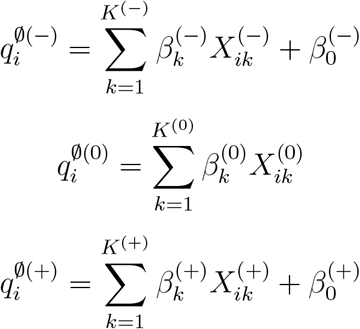

This is a standard multinomial logistic regression and has no hyperparameters that would necessitate the use of INLA. We approximate the posterior distribution over the regression coefficients using the same optimization and Laplace approximations used to evaluate the conditional posterior for the full model. We then draw a posterior sample from the reduced model, 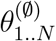, of the same size as the posterior sample obtained for the full model.

### Defining direct targets

We consider an exon a direct target if its splicing impact is sufficiently large to improve upon a reduced model without splicing activity. To assess this, we compare the posterior distribution of predictions between the full and the reduced models, *p* and *p*^∅^. In a knockdown experiment, enhancing targets should be downregulated and suppression targets upregulated. So we consider an exon an enhancing target if we are confident the splicing impact, *A*_*i*_, is negative enough to improve the downregulation prediction

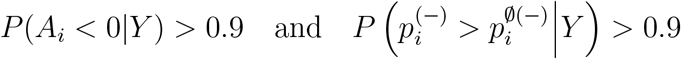

and a suppression target if the splicing impact is positive enough to improve the upregulation prediction

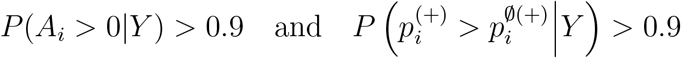

This assigns each exon into one of three mutually exclusive categories: enhancing target, suppression target, and nontarget. In some analyses, we wish to evaluate the confidence of these classifications and we do so with a multinomial regression that asks what fraction of exons with a given splicing impact were assigned to each category. This assigns each exon a probability of being an enhancing target, a nontarget, or a suppression target based on its splicing impact. We assumed that fractions of direct targets differ among upregulated, downregulated, and nonsignificant exons by fitting separate intercepts for each category.

### Evaluating eCLIP enrichment at targets

For SFs with significant activity maps and available eCLIP data, we downloaded the bed files representing peak calls for both replicates. We did not use the results of the irreproducible discovery rate analysis, because we believe this to represent an overly stringent criteria for combining replicate information. We considered the presence of a peak *>* 2-fold enriched over input in at least one of the eCLIP replicates as evidence of potential binding. The signal of differential splicing and eCLIP binding are clearest in highly expressed transcripts due to better representation in the sequencing reads; evaluating enrichment between differentially spliced and nonsignificant exons risks spurious associations due to correlated statistical power. To avoid this, we compared binding at targets to nontargets that significantly changed in the same direction. We further focused on the positions predicted by KATMAP to impact splicing. For both targets and nontargets, we counted the number of exons with evidence of binding, *k*_*target*_ and *k*_*nontarget*_, and used a beta-binomial conjugate model with a uniform prior to obtain 2,000 posterior samples of the fractions of bound target and nontarget exons, *f*_*target*_ and *f*_*nontarget*_

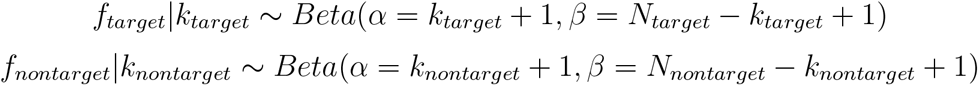

where *N*_*target*_ and *N*_*nontarget*_ are the total number of targets and nontargets that significantly changed in a given direction. We constructed a posterior sample of enrichment estimates as

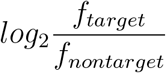

and computed the means and credible intervals from this.

### Defining effect sizes for changes in splicing

Differential splicing analyses typically report inclusion changes as Δ*ψ* = *ψ*_*kd*_ −*ψ*_*ctrl*_. However, under the hood, the statistical analysis is typically performed with respect to the log odds transformation of inclusion, 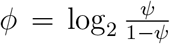, with the effect size being the log-odds ratio of inclusion before and after knockdown.

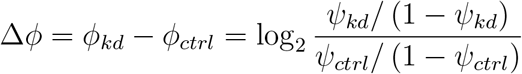

While our model does not aim to predict the magnitude of inclusion changes, only the direction, our analyses make use of inclusion-levels in two ways. First, we use the pre-knockdown inclusion level, *ϕ*_*ctrl*_ as a linear predictor, being informative of whether an increase or decrease in inclusion is detectable. Second, we assess whether our splicing impacts can predict the magnitudes of inclusion changes, despite not being trained on them. In both cases, we favor the log-odds representation.

The log-odds representations, *ϕ* and Δ*ϕ*, offer two advantages. First, *ψ* by definition must be between 0 and 1, and so the magnitude of Δ*ψ* only has meaning in the context of the original inclusion-level. Δ*ψ* = +0.1 is valid for *ψ*_*ctrl*_ = 0.05 but not for *ψ*_*ctrl*_ = 0.95, because *ψ*_*kd*_ = 1.05 is impossible. A second, related concern, is that inclusion levels near 0 or 1 likely represent more extreme splicing contexts than does intermediate inclusion. A change in inclusion from 0.5 to 0.6 corresponds to Δ*ψ* = 0.1, as does a change from 0.01 to 0.11. However, an exon included in only 1% of transcripts likely represents a strong silencing context or has weak splice sites, and so the increase of 10% likely reflects a more substantial change in regulation than a change from 50% to 60%. The odds-ratio reflects this, treating an inclusion change of 0.5 → 0.6 as representing a smaller effect (Δ*ϕ* ≈ +0.6) than a change of 0.01 → .11 (Δ*ϕ* ≈ +3.6).

### GO analysis of QKI-RBFOX cooperatively regulated exons

We defined confident QKI-RBFOX cotargets as predicted targets of either QKI or RBFOX that were downregulated upon knockdown and were inferred to also be regulated by the other factor based on splicing impacts. For the latter criteria, we chose a splicing impact cutoff of *<* −0.25 by examining the eCDFs (Fig. 5c,d). We constructed expression-matched background sets from all genes expressed in the QKI and RBFOX HepG2 knockdowns using importance sampling and used GOrilla to identify enriched GO terms relative to this background (Eden et al. 2009). We also performed this analysis using all differentially splicing QKI and RBFOX targets as the background. We used all three ontologies–process, function, and component. We identified significantly enriched terms using a 10% false discovery rate threshold.

### Inferring SF perturbations from splicing changes

To explain a set of splicing changes in terms of changes in SF regulation, we use the splicing impacts previously inferred from the knockdown data as the linear features in a multinomial regression. For each splicing factor, *f*, with a significant activity map, we use the posterior means of its activity 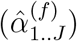 and binding parameters 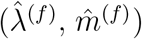 to compute splicing impacts for every considered exon *i*

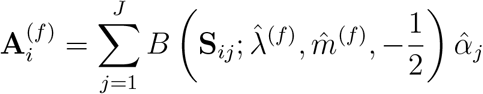

We incorporate these into a regression model, including linear coefficients to account for unperturbed inclusion-levels, expression, and their squares

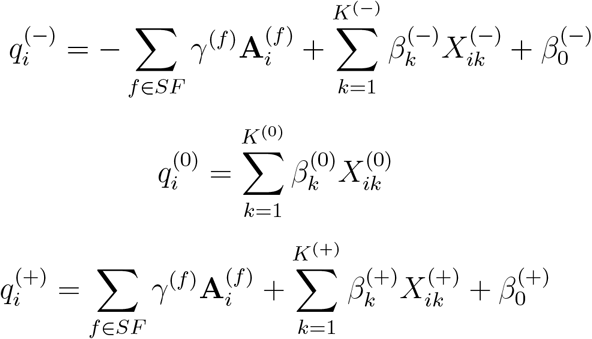

where each *γ*^(*f*)^ is a coefficient describing how predictive the splicing impacts for splicing factor *f* are of the splicing changes. And describe the observed splicing changes *Y*_1..*N*_ as following a categorical distribution

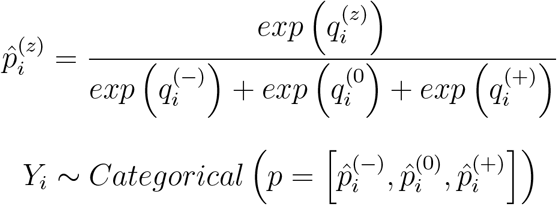

Because all of the splicing impacts **A** are precomputed this is a standard multinomial logistic regression. We use second-order optimization to find the posterior mode and, as described previously, use the curvature at the mode to approximate the posterior with a multivariate Gaussian. To assume sparse regression coefficients *post hoc*, we use ashr to apply shrinkage to the estimates.

In our analyses of the ENCODE data, we employ the same training data to define the observations and linear predictors used to learn the KATMAP models. When applying this approach the PDA cancer data, we used Supplemental Table 2 from (**empty citation**) to define the training data and sought to match the criteria used in the original analysis to define significant exons. We defined inclusion changes as metastatic versus primary, and restricted our analysis to cassette exons observed in at least 75% of samples, and defined exons as significantly up- or downregulated if inclusion changed by at least *±*10% and the reported FDR-corrected p-values was less than 0.05. We included all significant exons and 5,000 randomly selected nonsignificant exons in the training set. Because raw reads counts were not reported, we could not include log read counts as a linear feature, but include the average inclusion-level (in log-odds scale) from the primary tumor samples as a linear and quadratic predictor.

### Simulation study of model inference

For our simulation studies, we wished to assess how reliably our inference algorithm could recover the parameters which truly generated observed splicing changes. For our true parameters, (*θ, φ*)^(*true*)^,we used the posterior means inferred from twenty knockdowns that yielded significant activity maps. These parameters coupled with the predictor variables (*S* and *X*) naturally assign each exon probabilities of being downregulated, unchanged, or upregulated after knockdown. So that our simulated data provides enough statistical power to make confident inferences, we adjust the intercepts to ensure around 500 up- and downregulated exons. To generate the simulated observations *Y* ^(*sim*)^, we sample an outcome for each exon from a categorical distribution based on these predicted probabilities. We then use KATMAP to predict these simulated observations using the original predictor variables *S* and *X* to obtain a posterior sample (*θ, φ*)_1_, .., (*θ, φ*)_*N*_ |*Y* ^(*sim*)^. We computed posterior means and credible intervals and compared these to the true values (*θ, φ*)^(*true*)^.

### Statistical analyses

We represent uncertainty in our analyses with credible intervals. For KATMAP’s inferences, these are obtained by Monte Carlo sampling. With estimates based on count or presence/absence data, we employ the beta-binomial conjugate model, with uniform priors. For frequency estimates so obtained, we compute the mean and CI from the posterior beta distribution. For enrichment estimates, we sample frequencies from the posteriors of fore-ground and background to obtain a posterior sample of enrichments and construct the CI based on this. Otherwise, we typically employed Bayesian bootstrapping, wherein a posterior sample of estimates is obtained by repeatedly computing weighted averages, with the weights sampled from a uniform Dirichlet distribution (D. B. Rubin 1981).

There are several technical details consistent throughout our approach to inference. We use the jax scientific library to compute the gradients and Hessian matrices of our probability densities, which we need to optimize and quantify uncertainty in our model. We always work with log-probabilities for the sake of numerical stability. Despite this, loss of numerical precision occasionally results in non-positive semidefinite covariance matrices, that is matrices with eigenvalues less than zero. In this case, we follow theorem 3.2 in Cheng and Hingham (1998) to identify the nearest positive semidefinite matrix to Σ with eigenvalues greater than 10^−5^, which amounts to setting all eigenvalues less than 10^−5^ to this cutoff.

### Predicting the effects of deletions and splice-switching ASOs

To evaluate the effects of deletions and ASOs, we computed predictions for modified sequences. We assume that an ASO blocks SF interactions with the positions bound as well as one base on either side, and treat these positions as if they were the lowest affinity nucleotide when scoring with the binding model. For deletions, we simply score the sequence with the deletion. We use the posterior mean binding and activity parameters to construct our predictions. Because we wanted predict how each ASO or deletion alters regulation, rather than compute splicing impacts as

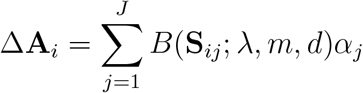

we instead computed predicted regulation for a given SF based on occupancy

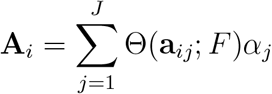

where **a**_*ij*_ = LogSumExp(*λ***S**_*ij*_) is the total affinity in bin *j*. We compute the change in regulation due to deletion/ASO *i* as

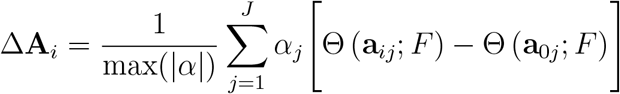

where **a**_0*j*_ is the affinity of region *j* in the reference sequence and **a**_*ij*_ is the affinity given the deletion/ASO. To compare changes in regulation across models, we normalize the predictions by the maximum change in regulation that could result from a complete loss of occupancy at a binding site (i.e. max(|*α*|). Doing this requires assuming a free protein concentration, *F*, which is not known *a priori*. As a proxy, we use estimates of free protein concentration in the cells from which the models were estimated based on the learned most-relevant-affinity parameter, *m*, and assuming free protein decreased by half after knockdown

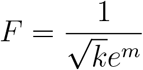

With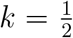. Our conclusions are not sensitive to choice of *k*, with 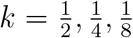 all predicting similar disruptions to regulation.

### Reporter assay

#### Constructing mutant sequences

For each selected exon, we identified the 10nt region most strongly contributing to the splicing impact (Fig. 4a,b). We then treated all regions that were at least 1/2 as impactful as potential SREs. For each SRE, we identified the three single-nucleotide polymorphisms most disruptive to binding. For exons with both up and downstream SREs, we constructed separate sequences with the upstream or downstream mutations alone, as well as with both upstream and downstream mutations.

#### Cell culture

HepG2 cells (ATCC) were grown in Dulbecco’s Modified Eagle Medium (GIBCO) supplemented with 10% bovine growth serum (GIBCO) and 1X penicillin-streptomycin (Gen Clone, Cat no. 25-512) in a 37°C incubator under 5% CO_2_.

#### Minigene splicing reporter constructs

RHCglo minigene reporter plasmid DNA (Addgene plasmid # 80169) was digested with SalI-HF (NEB, cat no. R3138S) and XbaI (NEB, cat no. R0145S) restriction enzymes to remove synthetic exon and flanking human *β*-globin intronic and splice site sequence. Digests were performed in a single reaction according to the manufacturer’s instructions, then purified using DNA Clean & Concentrator Kit (Zymo, Cat no. D4003). DNA constructs consisting of target exonic sequences + 250 nucleotides of flanking upstream and downstream intronic sequence, and sequence overlapping RHCglo sequence (Supplementary Table 5) were synthesized by Twist Bioscience and inserted into the linearized RHCglo pDNA using NEBuilder^®^ HiFi DNA Assembly Master Mix (NEB, cat no. E2621S). Reactions were used to transform Mix N Go DH5*α* cells (Zymo). Plasmid DNA from successful transformants were purified using Qiagen Spin Miniprep kit (Qiagen, cat no. 27104). RHCglo was a gift from Thomas Cooper.

#### Splicing factor knockdown and minigene experiments

For siRNA knockdown minigene reporter experiments 12-well plates were seeded 24h in advance with 0.1 × 10^6^ HepG2 cells. Cells were treated for 48h with 50 pmol of siRNA pool targeting the SF of interest (Supplementary Table 5) or ON-TARGETplus™ siRNA control pool using RNAiMAX reagent (Dharmacon™). At 48h media was removed and cells were transfected with 50 ng of minigene reporter pDNA using Lipofectamine 3000 reagent and incubated for 24h to allow transcript expression. Culture media was removed and cells were washed twice with 1X PBS pH 7.4 and total RNA was extracted using RNeasy`®` Mini kit (Cat no. 74106, Qiagen) according to the manufacturer’s instructions. Each experiment was performed as three biological replicates. The PCPB1 upstream mutant reporter assay was performed without the siRNA knockdown step as a single replicate and not used in later experiments due to the impact of mutations on the 3SS strength.

#### Minigene splicing quantification by semi-quantitative RT-PCR

First strand cDNA was generated from 100-150ng total extracted RNA using an Oligo d(T)_20_ primer (Invitrogen, cat no. 18418020) and SuperScript™ IV reverse transcriptase (Invitrogen, cat no. 18090050) according to the manufacturer’s instructions. cDNA was PCR amplified using RHCglo gene product specific primers RSV5U and RTRHC (universal to all constructs) and Q5^®^ High-Fidelity 2x Master Mix (NEB, cat no. M0492S) for 28 cycles. PCR products were resolved on 3% agarose gels and band intensity for exon inclusion and exon skipping isoforms was quantified using the ImageJ software package. PSI was calculated as band intensity of the inclusion isoform divided by total intensity of both isoforms.

#### Confirmation of SF knockdown by Western blot

HepG2 cells were treated with each siRNA and minigene combination as described above. Cells were washed with 1 x PBS then lysed in RIPA buffer (50 mM Tris (pH 7.0), 150 mM NaCl, 0.1% SDS, 0.5% sodium deoxycholate, 1X Triton x-100). Protein from each lysate was resolved on Bis-Tris NuPAGE acrylamide gels (Invitrogen) then transfered to a iBlot`®` nitrocellulose membrane (Cat no. IB4301001) using the iBLOT`®` dry blot system. Membranes were blocked for 1h in Azure Fluorescent Blot Blocking Buffer (Azure Biosystems Inc). Rabbit primary antibodies for each SF (Supplementary Table 5) at 1:1000 dilution were added to the membranes and incubated at 4°C overnight with gentle agitation. Membranes were washed with 1X TBST then incubated with fluorescently labeled secondary antibody (see Supplementary Table 5) at a 1:10,000 dilution for 1h at room temperature, washed with 1X TBST and visualized using Azure c600 (Azure Biosystems Inc). Each membrane was then incubated with mouse anti-*α*-tubulin (1:2500) for 1 hour at room temperature, washed with 1 X TBST, and incubated with secondary antibody (Supplementary Table 5) at room temperature for 1 hour, washed and visualized. Band intensity was quantified using ImageJ software and KD efficiency was calculated as *α*-tubulin normalized intensity of the SF’s band in the SF siRNA treated cells over the control siRNA treated cells (Supplementary Table 5).

#### Computational resources

All analyses were performed on a Dell Precision 7740 laptop with an NVIDIA Quadro RTX 3000 GPU (5.9GiB, CUDA version 11.4), 16 CPUs (Intel(R) Xeon(R) E-2286M, 2.40GHz), and 67.2 GB of RAM plus 137.4 GiB swap.

#### Software availability

KATMAP is available at https://gitlab.com/LaptopBiologist/katmap

## Acknowledgments

We thank members of the Burge lab for helpful discussions. This work was supported by an NIH F31 Fellowship (to M.P.M.) and by NIH grants (to C.B.B.).

## Author contributions

M.P.M. and C.B.B conceived the study. M.P.M. designed the study, developed the model and software, and processed, analyzed, and visualized the data. M.P.M, D.C.M., and C.B.B. designed the experiments and D.C.M. performed the experiments. M.P.M. wrote the original draft; all authors contributed to manuscript editing and support the conclusions

## Supplementary Material

**Supplementary Data 1:** A compressed archive containing KATMAP outputs for the activity maps used in our analyses. This includes the configuration files, training data, and motif models used to run each analysis; visual summaries of inferred activity and binding; the joint posterior samples over all model parameters; and the tables describing the inferred direct targets of each factor.

**Supplementary Table 1:** High-level overview of the KATMAP analyses. For each analysis performed, the source and accession of the differential splicing results and the motif models are provided, as well as descriptions of exons used to train the model. High-level summaries of the results are reported as the model comparison z-scores and numbers of significant activity coefficients, along the average number of predicted binding changes per exon. Whether an analysis yielded a significant activity map is indicated for each analysis, and if a given SF had both an RBNS-based PSAM and a PWM we indicate which model we used in our analyses.

**Supplementary Table 2:** Splicing impacts computed for all cassette exons in the ENCODE knockdowns for each SF with a significant activity map. Each row describes an exon, with the chromosomal coordinates denoting the first and last nucleotide of the exon, along with the splicing impacts assigned to that exon from KATMAP’s activity models for the different SFs. If an SF gave a significant activity model in more than one knockdown, we include the splicing impacts from the model with strongest evidence of splicing activity. **Supplementary Table 3:** Summary of exons identified as being coregulated by QKI and RBFOX based on the QKI (HepG2) and RBFOX2 (HepG2) knockdowns and the mouse *Rbfox* 1/2/3 triple knockdown data.

**Supplementary Table 4:** Splicing factor perturbations inferred from the ENCODE RNA-binding protein (RBP) knockdown datasets (including non-SF RBPs) using KATMAP’s activity models. Each row represents the results of a multinomial regression analysis using precomputed splicing impacts as predictors. The optimal coefficients, associated standard deviations, and sparse shrinkage estimates are provided in separate sheets.

**Supplementary Table 5:** Details on minigene splicing assays. Contains sequences for semiquantitative RT-PCR primers. Also includes siRNAs, primary and secondary antibodies used in minigene reporter experiments. Final sheet contains the name and measured knockdown efficiency for each RBP/minigene combination normalized to *α*-tubulin, as well as coordinates (hg19) and exonic/intronic sequences used to construct minigene reporters.

## Extended methods

### Adaptive importance sampling from the marginal posterior

Before describing the details of this procedure, it is important to note that we work with unconstrained representations of the hyperparameters. That is, for the length-scale *L* and stationary scale *σ*, we use their log-transformed values. If we denote the hyperparameters *φ*_*i*_ = (*m*_*i*_, *λ*_*i*_, *L*_*i*_, *σ*_*i*_), then we denote unconstrained version as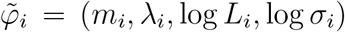. Computing the probability density on the unconstrained hyperparameters requires accounting for how the transformation stretches or compresses the density, so 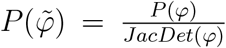 where *JacDet* is the Jacobian determinant of the transform. In practice, this amounts to multiplying the constrained density by *L*_*i*_ × *σ*_*i*_.

We begin using unconstrained version of the prior as our first proposal distribution

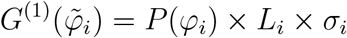

From this, we sample *n* unconstrained hyperparameters, 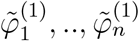 and compute the importance weights which describe how the proposal distribution differs from the target marginal posterior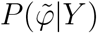:

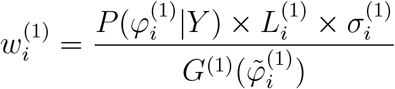

We use these evaluation points to estimate a new proposal distribution, *G*^(2)^ that better matches the target posterior, with an adaptation procedure rooted in fitting a Student’s T mixture model (TMM) with importance weighted expectation-maximization (IW-EM), described below. From this, we generate and evaluate a new sample of hyperparameters. We repeat this procedure, obtaining a series of proposal distributions that approach the target, until we are confident that we will draw a sufficiently large sample from the target.

At each iteration *τ*, this yields a set of evaluation points drawn from different proposal distributions

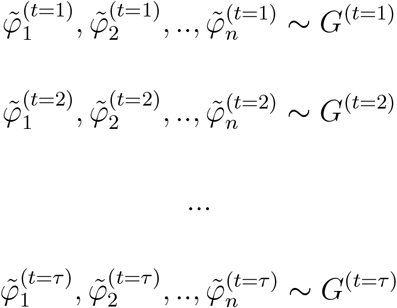

Each proposal distribution contributed equally to the set of generated evaluation points, so we compute the proposal probabilities from an equally weighted mixture of all past proposal distributions

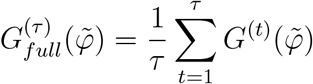

Then the raw importance weights are

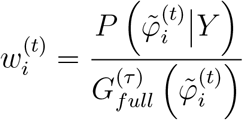

and the raw acceptance probabilities are

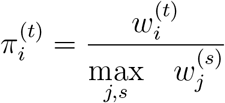

Following Vehtari et al, we apply a stabilizing correction to these importance weights, which we describe below, to obtain 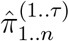

From these acceptance probabilities, we can determine what fraction of samples we expect to accept if we were to stop at the current iteration

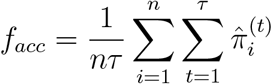

which also provides an estimate of the effective sample size

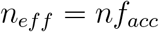

To determine whether we can stop iterating, we ask whether there is at least a 99% probability the number of accepted samples *n*_*acc*_ exceeds the desired sample size *n*_*final*_. The number of samples we will accept,*n*_*acc*_, is the sum of independent Bernoulli trials with variable success probabilities and so follows a Poisson-binomial distribution. As this distribution is never more variable than a binomial distribution with the same mean and number of trials, we use the binomial distribution to obtain a conservative estimate of the lower bound on *n*_*acc*_.

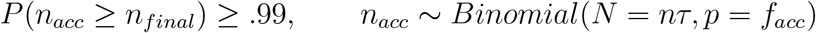

### Adapting the proposal distribution

We first use the EM procedure described below to fit a Student’s T mixture model (TMM) to the target, obtaining mixing weights *ω*_1..*C*_, means *µ*_1..*C*_, and covariance matrices Σ_1..*C*_, as well as a degree-of-freedom parameter *ν* shared by all *C* component distributions. This provides an estimate of the target that can account for both its local structure and the heaviness of its tails. However, in the first few iterations of adaptive importance sampling the effective sample is typically small (*<* 5) and we found the TMM-based proposal to occasionally become too concentrated around the modes of the target density, requiring extra rounds of adaption to properly explore tails. To decrease the time to convergence, we employ two heuristics to stabilize the adaptation in the early rounds of importance sampling. Both heuristics aim to ensure the proposal is distributed around the target density, but always somewhat more dispersed than the true target.

First, we double the fitted the covariance matrices, defining the mixture distribution which captures the local structure of the target density as

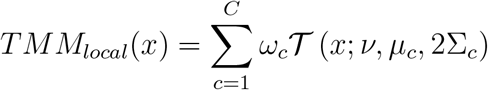

By intentionally making the proposal broader than the IW-EM estimate of the target density, we sacrifice somewhat the efficiency with which we obtain samples from the target density. However, this ensures that subsequent evaluation points fall in the tails of the target, decreasing the risk that the adaptation overshoots.

However, when the effective sample from the target is small, we lack precise information about the finer details of the distribution’s shape and *TMM*_*local*_ may still overfit. We therefore construct an even more conservative proposal. When *n*_*eff*_ is close to zero, IW-EM will shift the means in the direction of a mode in the target, but will learn little about the target’s covariance. Given that covariance of the target remains obscure, we may as well continue to use the previous proposal distribution. We approximate this by computing the covariance of the proposals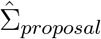. If *n*_*eff*_ is somewhat larger than zero, we might have enough information to estimate the global covariance of the target, 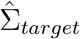, which we can obtain by computing an importance weighted covariance from the proposals. To find a intermediate between these two extremes, we take a weighted average that approaches 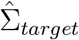as *n*_*eff*_ increases

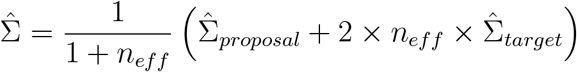

The Bayesian flavor of transitioning from an agnostic *a priori* assumption to a data-driven estimate as more information becomes available is not coincidental. This estimator can be thought of as being twice the posterior mean, Σ^*′*^, of the target covariance estimated using an inverse-Wishart Gaussian conjugate model with a prior scale matrix based on the initial proposal distribution, 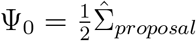, and *ν*_0_ = *p* + 2 degrees of freedom, where *p* is the number of rows/columns in the covariance matrix,

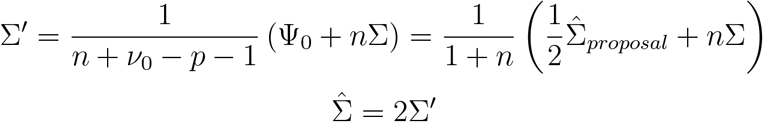

where Σ is a covariance matrix computed from *n* data points.

From this, we construct our more conservative proposal distribution

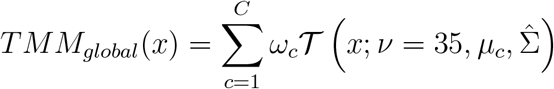

This uses the IW-EM estimates of target’s location, but replaces the covariances with a global estimate of the target’s covariance, to ensure new proposals explore the target density’s tails.

Our final proposal overlays these as a mixture, with the mixing weights depending upon *n*_*eff*_

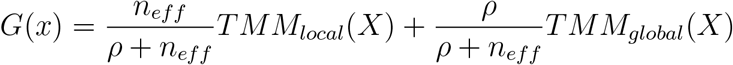

As *n*_*eff*_ increases, the composition of the mixture shifts from the conservative *TMM*_*global*_ to *TMM*_*local*_. We choose *ρ* = 4 such that when *n*_*eff*_ = 1, 80% of the new proposals come from the conservative mixture, at *n*_*eff*_ = 20 it is about 16% proposals, and when *n*_*eff*_ = 100 the proposal is largely derived from the local TMM, with less than 4% of proposal derived from the conservative mixture. Whatever *n*_*eff*_, the updated proposal always moves towards modes of the target density, because both *TMM*_*local*_ and *TMM*_*global*_ use the means and mixing proportions estimated from IW-EM.

### Importance-weighted Expectation Maximization for a multivariate T-mixture

To construct proposals that can robustly adapt to the target, we follow Cappé et al. (2008) and fit a TMM by importance weighted Expectation-Maximization (IW-EM). One approach to conceptualizating a multivariate T distribution is a scale-mixture of multivariate Gaussians. So a mixture of Student T distributions can be described as a mixture of Gaussians with two latent variable per realization *x*_*i*_

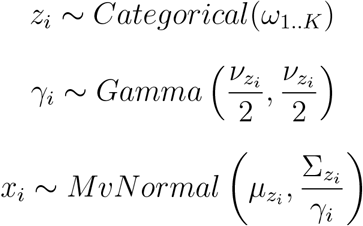

where *ν* is the degrees of freedom. *ω*_1..*K*_ is a vector of mixing weights describing the fraction of the full mixture each component distribution contributes. *µ*_1..*K*_ and Σ_1..*K*_ are means and covariances of the *K* component distributions. *z*_1..*N*_ is vector of unobserved variables indicating which component each observation *x*_*i*_ truly derives from. The second set of unobserved variables *γ*_1..*N*_ come from the scale-mixture representation of the Student T distribution and describe how Gaussian *z*_*i*_ was rescaled to generate each realization *x*_*i*_.

While, Cappé et al. (2008) initialize once and perform a single EM-update at each step of importance sampling, we instead reinitialize at each importance sampling iteration *t* and perform *t* EM-updates to prevent the adaptive distribution from getting trapped. We use K-means clustering of the proposals with 10 × *t* components, to define the initial means ensure the components are well-distributed throughout the proposal distribution.

### Expectation Step

For each observed *x*_*i*_, there are two unobserved variables, *z*_1..*N*_ and *γ*_1..*N*_. The expectation step infers these latent variables by computing their expected values given the parameters of the mixture distribution (*ω*_1..*K*_, *µ*_1..*K*_, Σ_1..*K*_).

To estimate the probability that *x*_*i*_ arose from a given component distribution *d*, we compute

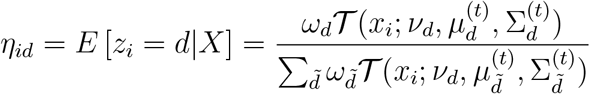

where 𝒯 denotes the multivariate Student-t probability density function as implemented in scipy.

Then to estimate the scale of the Gaussian from which *x*_*i*_ arose, assuming it belongs to the *d*-th Student T distribution, we compute

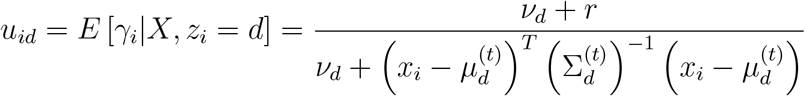

where *r* is the dimension of the distribution (in our case, the number of hyperparameters in the additive model).

### Maximization Step

The maximization step uses these estimates of the latent variables to refine the estimates of the mixture distribution’s parameters. These estimates are essentially averages weighted by the component probabilities *η*_*id*_. To fit the mixture to the target rather than the proposals, we follow Cappé et al. (2008) compute the updates as weighted averages using the importance weights *π*_*i*..*N*_ to weight the observations:

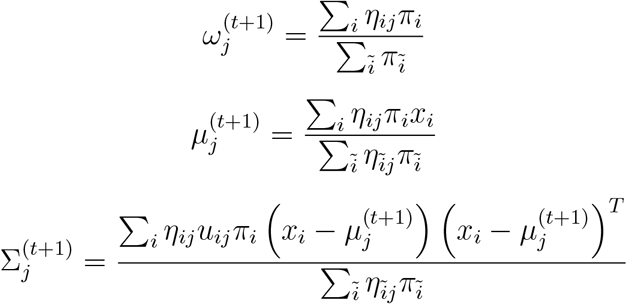

The degree-of-freedom parameter, *ν*, controls the tails of the distribution. There is no closed-form estimate of *ν* and so it must be estimated by optimization. To make this tractable, we constrain the components of the mixture to all share the same *ν* parameter. By reducing this to a single parameter, we can use a binary search to update *ν*^(*t*+1)^, considering values between *ν* ∈ (4, 100). The importance weighted EM scheme optimizes the entropy of the normalized importance weights we would get if we used the mixture as the proposal distribution

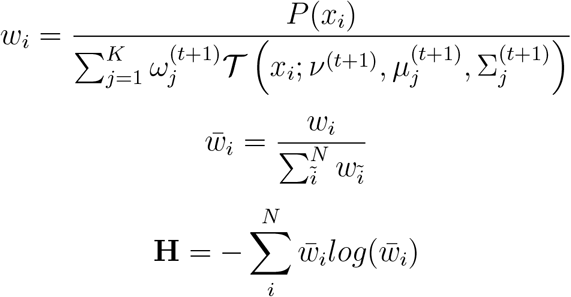

So we evaluate the entropy at twenty values of *ν* between 4 and 100 and identify all local maxima on this grid. We then perform binary searches between the neighbors of each maxima to hone in on the true local maxima and set *ν*^(*t*+1)^ to the value of *ν* that yielded the highest entropy.

### Stabilizing the importance weights

The set of importance weights 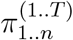 represent an estimate of how the proposal distribution differs from the target distribution. To properly characterize the target distribution, the importance weights must be based on a sufficient number of proposals spanning typical set of the target. If the proposal distribution is very different from the target, the computed importance weights will poorly characterize the target and importance sampling will be inefficient or untrustworthy.

Vehtari et al provide an approach for obtaining more stable estimates of the importance weights by fitting a Pareto distribution to the tail of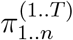. Using the implementation of this Pareto smoothing from the arviz library (Kumar et al. 2019), we compute smoothed weights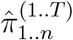. Importantly, the shape parameter of the fitted Pareto distribution, 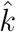, is diagnostic of whether importance sampling will be reliable.

We only accept posterior samples based on the Pareto-smoothed importance weights. Occasionally in the early iterations of adapting the proposal distribution, however, the range of values computed for the importance weights may be too extreme for Pareto smoothing to yield reliable importance estimates (e.g.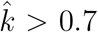). In these cases, we use an older, simpler approach to stabilize the weights (Ionides 2008). This defines a cutoff based on the total number of importance weights *N*_*π*_ and average importance weight 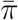

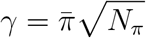

and then replaces all importance weights greater than *γ* with *γ*

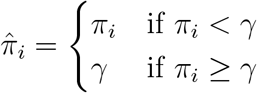

The way in which this flattens the importance weights will bias the importance estimates by accepting some proposals which should be rejected. When used to adapt the proposal distribution, this should result in an estimate of the target distribution that is broader than the true target. This decreases the efficiency with which the proposal is adapted, but prevents it from collapsing to a subset of the target distribution.

### Assumptions about the spatial correlations: *L, σ*

There are two identifiability issues that can arise with Gaussian process priors. If the length-scale *L* is near zero, it corresponds to the biologically implausible assumption of complete spatial independence of the activity coefficients. But it also introduces problems for statistical inference. As we perform inference with respect to the log-transform of *L*, all values of *logL* ≪ 0 are essentially equivalent. Similarly, length-scales much larger than the maximum distance between regions analyzed *L* ≫ *max*(|*j* − *k*|) are practically indistinguishable.

To prevent the inference algorithm from wasting computation by exploring large but practically equivalent regions of parameter space, we construct a prior that avoids both large and small values of *L*. To avoid overly large length-scales, we use the right-tail of a generalized gamma distribution

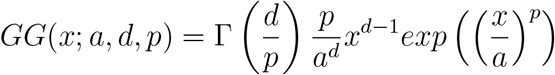

This does not prevent *L* from being close to zero. We avoid these small values by convolving the generalized gamma with the left tail of an inverse-gamma distribution

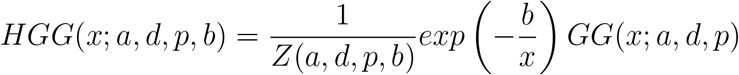

where the normalizing constant is computed numerically (‘scipy.integrate.quad’)

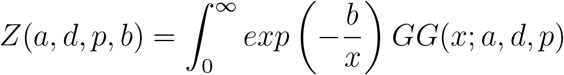

We set the prior on the length-scale as

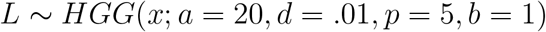

which places 99% of the prior probability on the interval 0.32 *< L <* 22.6.

We use this same distribution to exclude small values of the stationary standard deviation. If *σ* is close to zero, all of the activity coefficients become constrained to be close to zero solely by the prior. One way of conceptualizing the prior on *σ* is that it reflects different assumptions we might make about the regression coefficients in our regression analysis. We would never perform a regression analysis where we assumed *a priori* that all of the regression coefficient were zero. Whether the coefficients should be close to zero is something we wish to learn from the data, not assume in advance. So we define our prior on the stationary standard deviation as

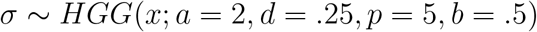

which places 99% of the prior probability on the interval 0.14 *< σ <* 8.7

### Model comparison against the reduced model

To evaluate whether the data support splicing activity, we compare the full model against a reduced model without splicing activity. To verify that any improvement over the reduced model did not reflect overfitting to the training data, we sought to evaluate each model’s performance on held-out data. However, to sidestep the need to rerun the inference algorithm dozens of times for each knockdown, we employ the Pareto-smoothed importance sampling approximation to leave-one-out (PSIS-LOO) cross-validation proposed by Vehtari, Simpson, et al. (2024)

PSIS-LOO uses the posterior sample from training on all data as a proposal distribution to obtain estimates regarding the leave-one-out posterior distribution that would be obtained by excluding the i-th observation. It leverages the insight that the ratio between the complete and the leave-one-out posteriors depends only on the left-out observation, with all other terms in the posteriors cancelling out.

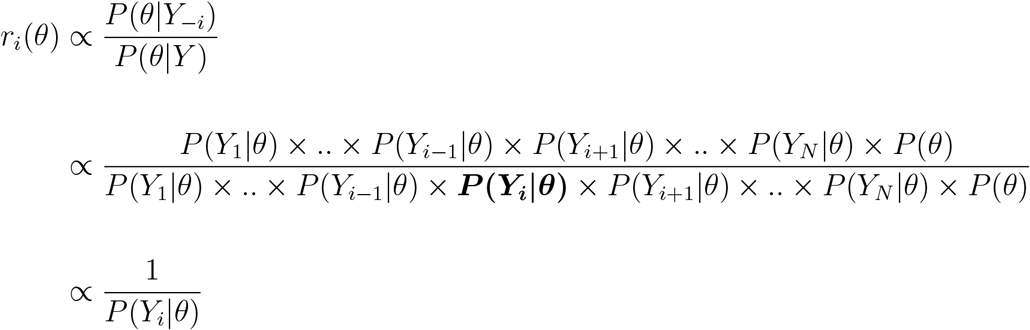

These importance weights are then stabilized using the aforementioned Pareto smoothing to obtain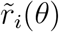. We estimate the log expected posterior predictive probability for each held-out exon *i* by computing the importance-weighted average over the complete posterior sample

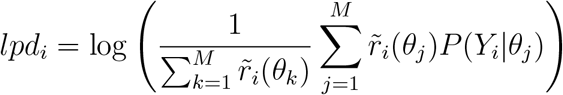

To combine this across all held-out exons, we sum the estimated log-probabilities, exactly as we would when computing the model’s objective function

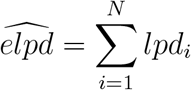

We chose this estimator as it is the finite sample version of the widely-applicable information criterion (WAIC). We compute this for the full, 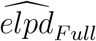, and reduced models, 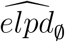, and summarize the model comparison as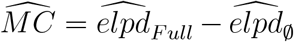. A positive value of 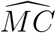 means the full model with splicing activity is a better explanation of the data than the reduced model without splicing activity. To quantify how confident we are in this evidence for splicing activity, we bootstrap the exons 10,000 times, estimating 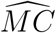 each time, and use these obtain a standard deviation *SD*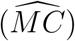. We then compute a Z-score

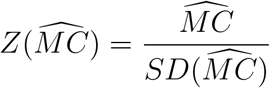

### Inferring the underlying correlation of variables with known uncertainty

In some of our analyses we wanted to determine the correlation between estimated values 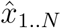 and ŷ_1..*N*_, when each of the estimates was associated with known uncertainty *σ*_*x*,1..*N*_ and *σ*_*y*,1..*N*_. Simply computing the correlation between 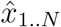 and ŷ_1..*N*_ would underestimate the true association between the true values *x* and *y*, which is obscured by the uncertainty in the estimates. We therefore employ a hierarchical model to disentangle the underlying association from the uncertainty in the individual estimates.

We first assume that the estimated values 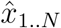 and ŷ_1..*N*_ are normally distributed around the true values *x*_1..*N*_ and *y*_1..*N*_ with standard deviations *σ*_*x*,1..*N*_ and *σ*_*y*,1..*N*_

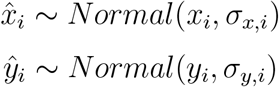

we then assume that the underlying true values follow a latent multivariate normal distribution with the correlation structure of interest

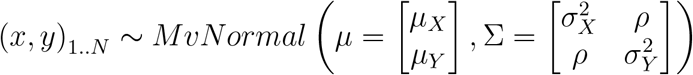

Our primary goal is infer *ρ*, disentangled from the uncertainty. For the full hierarchical model, we place priors on the means and standard deviations of this latent Gaussian using the means and standard deviations of the noisy estimates 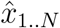 and ŷ_1..*N*_. We also place a relatively uninformative prior on the correlation coefficient in logit scale.

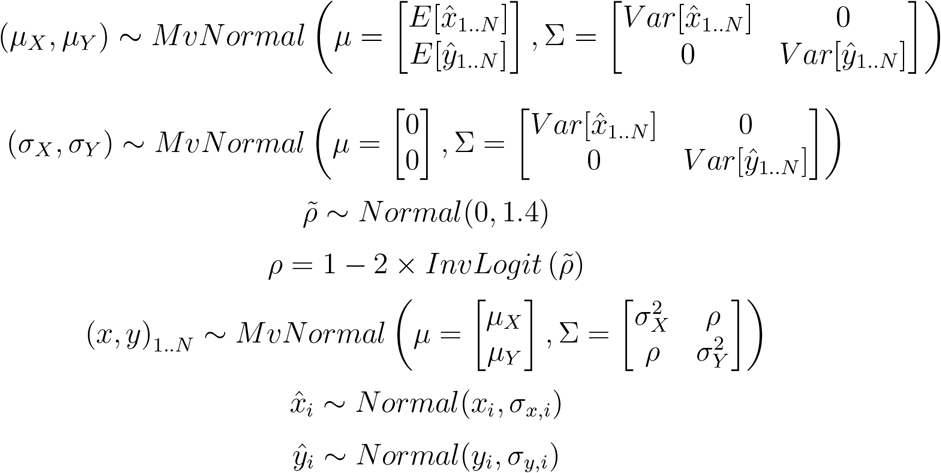

In practice, the parameters of hierarchical models often have complicated associations with each other that prevents efficient exploration of the posterior. We follow the standard practice to resolve this by using the noncentered parameterization of the latent Gaussian and representing (*x, y*)_1..*N*_ as the product of a unit normal distribution and the Cholesky factor of the covariance matrix

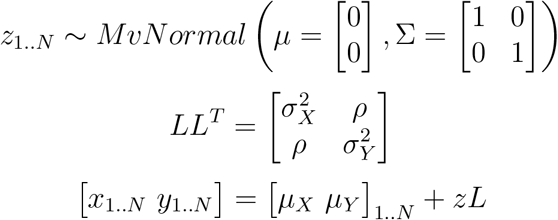

We use the No-U-Turn sampler, as implemented in PyMC3 (3.11.2), to draw 4,000 samples from the joint posterior over all parameters (*ρ, µ*_*X*_, *µ*_*Y*_, *σ*_*X*_, *σ*_*Y*_, *x*_1..*N*_, *y*_1..*N*_) in 4 chains with the target acceptance probability set to 0.9.

### Working with differential splicing analyses

#### Recovering effect sizes estimates from rMATS outputs

rMATs reports the inclusion levels and inclusion level difference using direct summaries of the read counts, rather than the effect sizes inferred by the underlying model. The actual parameters of the rMATS model, however, are defined in log-odds scale. This log-odds representation allows interval estimates to be naturally defined around the effect size in a way that respects the constraint that 0 *< ψ <* 1. The read count based summaries, however, may equal 0 or 1, and cannot be converted as is to log-odds scale. To resolve this, we recompute PSI with pseudocounts added to the observed reads, which is equivalent to estimating the inclusion read fraction with a beta-binomial model.

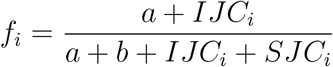

We identify the appropriate pseudocounts from the data itself, fitting a beta-binomial distribution to the read counts of all exons. We then estimate the inclusion read fraction for each exon and convert these to PSI

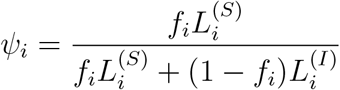

where the 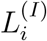 and 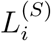 account for the differing amounts of informative sequence associated with the inclusion and exclusion junctions respectively.

### Reconstructing uncertainty about effect size from rMATS outputs

We sought to reconstruct the posterior standard deviation for these effect sizes from the rMATS p-values. rMATS computes its p-values using a likelihood ratio test, comparing the likelihood of the full model to that where the effect size is set to a small value near zero,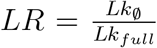. The test statistic Λ_*i*_ = −2*logLR* follows a chi-squared distribution and can be recovered by inputting the rMATS p-value, *p*_*i*_ for exon *i* into the ch-squared inverse cumulative density function

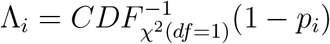

Describing the uncertainty about the effect sizes with a mean and standard deviation implies a normal approximation to the likelihood. The likelihood-ratios, therefore, can be thought of as comparing the density of this normal distribution at its mode to a point in its tail. The likelihood of the full model is evaluated at its mode (e.g. effect size)

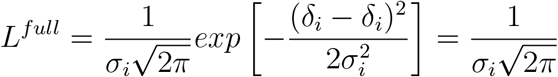

and is entirely determined by the standard deviation. The null model, however, evaluates the likelihood at a point away from the mode, *d*_∅_, assuming the effect size is close to zero

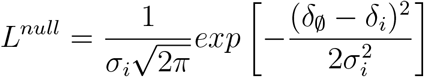

The likelihood ratio is then

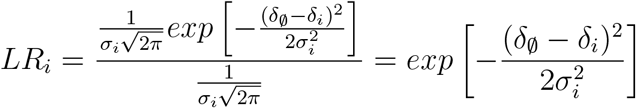

and the negative log-likelihood ratio is

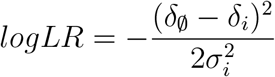

To get the variance 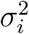 of the normal approximation we can rearrange

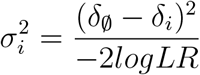

The Chi2 test statistic, Λ_*i*_ = −2*logLR*_*i*_, is the denominator so this can be written:

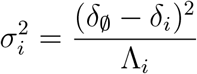

## References

Bak, Maciej et al. (May 15, 2024). “MAPP unravels frequent co-regulation of splicing and polyadenylation by RNA-binding proteins and their dysregulation in cancer”. In: Nature Communications 15.1. Publisher: Nature Publishing Group, p. 4110. issn: 2041-1723. doi: 10.1038/s41467-024-48046-1.

Balwierz, Piotr J. et al. (May 1, 2014). “ISMARA: automated modeling of genomic signals as a democracy of regulatory motifs”. In: Genome Research 24.5. Company: Cold Spring Harbor Laboratory Press Distributor: Cold Spring Harbor Laboratory Press Institution: Cold Spring Harbor Laboratory Press Label: Cold Spring Harbor Laboratory Press Publisher: Cold Spring Harbor Lab, pp. 869–884. issn: 1088-9051, 1549-5469. doi: 10.1101/gr.169508.113.

Best, Andrew et al. (Sept. 11, 2014). “Human Tra2 proteins jointly control a CHEK1 splicing switch among alternative and constitutive target exons”. In: Nature Communications 5, p. 4760. issn: 2041-1723. doi: 10.1038/ncomms5760.

Boutz, Paul L. et al. (July 1, 2007). “A post-transcriptional regulatory switch in polypyrimidine tract-binding proteins reprograms alternative splicing in developing neurons”. In: Genes & Development 21.13. Company: Cold Spring Harbor Laboratory Press Distributor: Cold Spring Harbor Laboratory Press Institution: Cold Spring Harbor Laboratory Press Label: Cold Spring Harbor Laboratory Press Publisher: Cold Spring Harbor Lab, pp. 1636–1652. issn: 0890-9369, 1549-5477. doi: 10.1101/gad.1558107.

Bradbury, James et al. (2018). JAX: composable transformations of Python+NumPy programs. Version 0.3.13.

Bradley, Robert K. and Olga Anczuków (Mar. 2023). “RNA splicing dysregulation and the hallmarks of cancer”. In: Nature Reviews. Cancer 23.3, pp. 135–155. issn: 1474-1768. doi: 10.1038/s41568-022-00541-7.

Brosseau, Jean-Philippe et al. (Feb. 1, 2014). “Tumor microenvironment–associated modifications of alternative splicing”. In: RNA 20.2. Company: Cold Spring Harbor Laboratory Press Distributor: Cold Spring Harbor Laboratory Press Institution: Cold Spring Harbor Laboratory Press Label: Cold Spring Harbor Laboratory Press Publisher: Cold Spring Harbor Lab, pp. 189–201. issn: 1355-8382, 1469-9001. doi: 10.1261/rna.042168.113.

Cao, Wenguang et al. (Sept. 2012). “Control of alternative splicing by forskolin through hnRNP K during neuronal differentiation”. In: Nucleic Acids Research 40.16, pp. 8059– 8071. issn: 0305-1048. doi: 10.1093/nar/gks504.

Cappè, Olivier et al. (Dec. 1, 2008). “Adaptive importance sampling in general mixture classes”. In: Statistics and Computing 18.4, pp. 447–459. issn: 1573-1375. doi: 10.1007/s11222-008-9059-x.

Chen, Charlie Degui, Ryuji Kobayashi, and David M. Helfman (Mar. 1, 1999). “Binding of hnRNP H to an exonic splicing silencer is involved in the regulation of alternative splicing of the rat -tropomyosin gene”. In: Genes & Development 13.5, pp. 593–606. issn: 0890-9369.

Chen, Xinyuan et al. (July 3, 2023). “The RNA-binding proteins hnRNP H and F regulate splicing of a MYC-dependent HRAS exon in prostate cancer cells”. In: Proceedings of the National Academy of Sciences of the United States of America 120.28, e2220190120. issn: 0027-8424. doi: 10.1073/pnas.2220190120.

Cheung, Hannah C. et al. (Aug. 1, 2009). “Splicing factors PTBP1 and PTBP2 promote proliferation and migration of glioma cell lines”. In: Brain 132.8, pp. 2277–2288. issn: 0006-8950. doi: 10.1093/brain/awp153.

Chou, Min-Yuan et al. (Jan. 1999). “hnRNP H Is a Component of a Splicing Enhancer Complex That Activates a c-src Alternative Exon in Neuronal Cells”. In: Molecular and Cellular Biology 19.1, pp. 69–77. issn: 0270-7306.

Coelho, Miguel B et al. (Mar. 4, 2015). “Nuclear matrix protein Matrin3 regulates alternative splicing and forms overlapping regulatory networks with PTB”. In: The EMBO Journal 34.5. Num Pages: 668 Publisher: John Wiley & Sons, Ltd, pp. 653–668. issn: 0261-4189. doi: 10.15252/embj.201489852.

Duttke, Sascha H. et al. (July 2024). “Position-dependent function of human sequencespecific transcription factors”. In: Nature 631.8022, pp. 891–898. issn: 1476-4687. doi: 10.1038/s41586-024-07662-z.

Eden, Eran et al. (Feb. 3, 2009). “GOrilla: a tool for discovery and visualization of enriched GO terms in ranked gene lists”. In: BMC Bioinformatics 10.1, p. 48. issn: 1471-2105. doi: 10.1186/1471-2105-10-48.

Ellingford, Jamie M. et al. (July 19, 2022). “Recommendations for clinical interpretation of variants found in non-coding regions of the genome”. In: Genome Medicine 14.1, p. 73. issn: 1756-994X. doi: 10.1186/s13073-022-01073-3.

Finkel, Richard S. et al. (Dec. 17, 2016). “Treatment of infantile-onset spinal muscular atrophy with nusinersen: a phase 2, open-label, dose-escalation study”. In: Lancet (London, England) 388.10063, pp. 3017–3026. issn: 1474-547X. doi: 10.1016/S0140-6736(16)31408-8.

Ghanem, Louis R. et al. (July 30, 2018). “Poly(C)-Binding Protein Pcbp2 Enables Differentiation of Definitive Erythropoiesis by Directing Functional Splicing of the Runx1 Transcript”. In: Molecular and Cellular Biology 38.16, e00175–18. issn: 0270-7306. doi: 10.1128/MCB.00175-18.

Graveley, Brenton R., Klemens J. Hertel, and Tom Maniatis (Nov. 16, 1998). “A systematic analysis of the factors that determine the strength of pre-mRNA splicing enhancers”. In: The EMBO Journal 17.22. Num Pages: 6756 Publisher: John Wiley & Sons, Ltd, pp. 6747–6756. issn: 0261-4189. doi: 10.1093/emboj/17.22.6747.

Guil, Sónia et al. (Apr. 2003). “Roles of hnRNP A1, SR Proteins, and p68 Helicase in c-H-ras Alternative Splicing Regulation”. In: Molecular and Cellular Biology 23.8, pp. 2927–2941. issn: 0270-7306. doi: 10.1128/MCB.23.8.2927-2941.2003.

Han, Areum et al. (Jan. 30, 2014). “De Novo Prediction of PTBP1 Binding and Splicing Targets Reveals Unexpected Features of Its RNA Recognition and Function”. In: PLOS Computational Biology 10.1. Publisher: Public Library of Science, e1003442. issn: 1553-7358. doi: 10.1371/journal.pcbi.1003442.

Hardy, R. J. (Oct. 1, 1998). “QKI expression is regulated during neuron-glial cell fate decisions”. In: Journal of Neuroscience Research 54.1, pp. 46–57. issn: 0360-4012. doi: 10.1002/(SICI)1097-4547(19981001)54:1<46::AID-JNR6>3.0.CO;2-H.

Heiner, Monika et al. (Jan. 1, 2010). “HnRNP L-mediated regulation of mammalian alternative splicing by interference with splice site recognition”. In: RNA Biology 7.1. Publisher: Taylor & Francis eprint: 10.4161/rna.7.1.10402, pp. 56–64. issn: 1547-6286. doi: 10.4161/rna.7.1.10402.

Hua, Yimin et al. (Mar. 13, 2007). “Enhancement of SMN2 Exon 7 Inclusion by Antisense Oligonucleotides Targeting the Exon”. In: PLOS Biology 5.4. Publisher: Public Library of Science, e73. issn: 1545-7885. doi: 10.1371/journal.pbio.0050073.

Hugh-White, Rupert (2021). “Analysing RNA-seq datasets to determine how pre-mRNA splicing is regulated by RNA binding proteins and cis-acting elements”. PhD thesis. King’s College London.

Huttlin, Edward L. et al. (May 27, 2021). “Dual proteome-scale networks reveal cell-specific remodeling of the human interactome”. In: Cell 184.11, 3022–3040.e28. issn: 0092-8674. doi: 10.1016/j.cell.2021.04.011.

Ionides, Edward L. (June 1, 2008). “Truncated Importance Sampling”. In: Journal of Computational and Graphical Statistics. Publisher: Taylor & Francis. doi: 10.1198/106186008X320456.

Jacko, Martin et al. (Feb. 21, 2018). “Rbfox Splicing Factors Promote Neuronal Maturation and Axon Initial Segment Assembly”. In: Neuron 97.4, 853–868.e6. issn: 1097-4199. doi: 10.1016/j.neuron.2018.01.020.

Jaganathan, Kishore et al. (Jan. 24, 2019). “Predicting Splicing from Primary Sequence with Deep Learning”. In: Cell 176.3, 535–548.e24. issn: 1097-4172. doi: 10.1016/j.cell.2018.12.015.

Jbara, Amina et al. (May 2023). “RBFOX2 modulates a metastatic signature of alternative splicing in pancreatic cancer”. In: Nature 617.7959. Publisher: Nature Publishing Group, pp. 147–153. issn: 1476-4687. doi:10.1038/s41586-023-05820-3.

Jens, Marvin et al. (Nov. 9, 2022). RBPamp: Quantitative Modeling of Protein-RNA Interactions in vitro Predicts in vivo Binding. Pages: 2022.11.08.515616 Section: New Results. doi: 10.1101/2022.11.08.515616.

Ji, Xinjun et al. (Mar. 18, 2016). “CP binding to a cytosine-rich subset of polypyrimidine tracts drives a novel pathway of cassette exon splicing in the mammalian transcriptome”. In: Nucleic Acids Research 44.5, pp. 2283–2297. issn: 0305-1048. doi: 10.1093/nar/gkw088.

Karni, Rotem et al. (Mar. 2007). “The gene encoding the splicing factor SF2/ASF is a protooncogene”. In: Nature Structural & Molecular Biology 14.3. Publisher: Nature Publishing Group, pp. 185–193. issn: 1545-9985. doi: 10.1038/nsmb1209.

Kràlovičovà, Jana et al. (July 6, 2018). “PUF60-activated exons uncover altered 3 splice-site selection by germline missense mutations in a single RRM”. In: Nucleic Acids Research 46.12, pp. 6166–6187. issn: 0305-1048. doi: 10.1093/nar/gky389.

Krismer, Konstantin et al. (Aug. 25, 2020). “Transite: A Computational Motif-Based Analysis Platform That Identifies RNA-Binding Proteins Modulating Changes in Gene Expression”. In: Cell Reports 32.8, p. 108064. issn: 2211-1247. doi: 10.1016/j.celrep.2020.108064.

Kumar, Ravin et al. (Jan. 15, 2019). “ArviZ a unified library for exploratory analysis of Bayesian models in Python”. In: Journal of Open Source Software 4.33, p. 1143. issn: 2475-9066. doi: 10.21105/joss.01143.

Lambert, Nicole et al. (June 5, 2014). “RNA Bind-n-Seq: quantitative assessment of the sequence and structural binding specificity of RNA binding proteins”. In: Molecular Cell 54.5, pp. 887–900. issn: 1097-4164. doi: 10.1016/j.molcel.2014.04.016.

Lang, Benjamin et al. (July 9, 2021). “Matrix-screening reveals a vast potential for direct protein-protein interactions among RNA binding proteins”. In: Nucleic Acids Research 49.12, pp. 6702–6721. issn: 0305-1048. doi: 10.1093/nar/gkab490.

Leclair, Nathan K. et al. (Nov. 19, 2020). “Poison Exon Splicing Regulates a Coordinated Network of SR Protein Expression during Differentiation and Tumorigenesis”. In: Molecular Cell 80.4, 648–665.e9. issn: 1097-2765. doi: 10.1016/j.molcel.2020.10.019.

Li, Ji et al. (July 30, 2018). “An alternative splicing switch in FLNB promotes the mesenchymal cell state in human breast cancer”. In: eLife 7, e37184. issn: 2050-084X. doi: 10.7554/eLife.37184.

Lim, Kian Huat et al. (July 5, 2011). “Using positional distribution to identify splicing elements and predict pre-mRNA processing defects in human genes”. In: Proceedings of the National Academy of Sciences 108.27, pp. 11093–11098. issn: 0027-8424, 1091-6490. doi: 10.1073/pnas.1101135108.

Llorian, Miriam et al. (Sept. 2010). “Position-dependent alternative splicing activity revealed by global profiling of alternative splicing events regulated by PTB”. In: Nature Structural & Molecular Biology 17.9. Publisher: Nature Publishing Group, pp. 1114–1123. issn: 1545-9985. doi: 10.1038/nsmb.1881.

Merendino, L. et al. (Dec. 16, 1999). “Inhibition of msl-2 splicing by Sex-lethal reveals interaction between U2AF35 and the 3’ splice site AG”. In: Nature 402.6763, pp. 838– 841. issn: 0028-0836. doi: 10.1038/45602.

Nocedal, Jorge and Stephen Wright (Dec. 11, 2006). Numerical Optimization. Google-Books-ID: VbHYoSyelFcC. Springer Science & Business Media. 686 pp. isbn: 978-0-387-40065-5.

North, Khrystyna et al. (July 2022). “Synthetic introns enable splicing factor mutation-dependent targeting of cancer cells”. In: Nature biotechnology 40.7, pp. 1103–1113. issn: 1087-0156. doi: 10.1038/s41587-022-01224-2.

Page-McCaw, P. S., K. Amonlirdviman, and P. A. Sharp (Dec. 1999). “PUF60: a novel U2AF65-related splicing activity”. In: RNA (New York, N.Y.) 5.12, pp. 1548–1560. issn: 1355-8382. doi: 10.1017/s1355838299991938.

Rasmussen, Carl Edward and Christopher K. I. Williams (2006). Gaussian processes for machine learning. Adaptive computation and machine learning. MIT Press. I-XVIII, 1-248. isbn: 0-262-18253-X.

Ray, Debashish, Hilal Kazan, Esther T. Chan, et al. (July 2009). “Rapid and systematic analysis of the RNA recognition specificities of RNA-binding proteins”. In: Nature Biotechnology 27.7. Publisher: Nature Publishing Group, pp. 667–670. issn: 1546-1696. doi: 10.1038/nbt.1550.

Ray, Debashish, Hilal Kazan, Kate B. Cook, et al. (July 2013). “A compendium of RNAbinding motifs for decoding gene regulation”. In: Nature 499.7457. Publisher: Nature Publishing Group, pp. 172–177. issn: 1476-4687. doi: 10.1038/nature12311.

Rothrock, Caryn R, Amy E House, and Kristen W Lynch (Aug. 3, 2005). “HnRNP L represses exon splicing via a regulated exonic splicing silencer”. In: The EMBO Journal 24.15. Num Pages: 2802 Publisher: John Wiley & Sons, Ltd, pp. 2792–2802. issn: 0261-4189. doi: 10.1038/sj.emboj.7600745.

Rubin, Donald B. (Jan. 1981). “The Bayesian Bootstrap”. In: The Annals of Statistics 9.1. Publisher: Institute of Mathematical Statistics, pp. 130–134. issn: 0090-5364, 2168-8966. doi: 10.1214/aos/1176345338.

Rubin, Jonathan D. et al. (June 2, 2021). “Transcription factor enrichment analysis (TFEA) quantifies the activity of multiple transcription factors from a single experiment”. In: Communications Biology 4.1. Publisher: Nature Publishing Group, pp. 1–15. issn: 2399-3642. doi: 10.1038/s42003-021-02153-7.

Rue, Håvard, Sara Martino, and Nicolas Chopin (2009). “Approximate Bayesian inference for latent Gaussian models by using integrated nested Laplace approximations”. In: Journal of the Royal Statistical Society: Series B (Statistical Methodology) 71.2. eprint: https://onlinelibrary.wiley.com/doi/pdf/10.1111/j.1467-9868.2008.00700.x, pp. 319–392. issn: 1467-9868. doi: 10.1111/j.1467-9868.2008.00700.x.

Shen, Shihao et al. (Dec. 23, 2014). “rMATS: Robust and flexible detection of differential alternative splicing from replicate RNA-Seq data”. In: Proceedings of the National Academy of Sciences 111.51. Publisher: Proceedings of the National Academy of Sciences, E5593– E5601. doi: 10.1073/pnas.1419161111.

Sherman, Larry et al. (Nov. 1997). “Interdomain binding mediates tumor growth suppression by the NF2 gene product”. In: Oncogene 15.20. Publisher: Nature Publishing Group, pp. 2505–2509. issn: 1476-5594. doi: 10.1038/sj.onc.1201418.

Spellman, Rachel, Miriam Llorian, and Christopher W.J. Smith (Aug. 3, 2007). “Cross-regulation and Functional Redundancy between the Splicing Regulator PTB and Its Paralogs nPTB and ROD1”. In: Molecular Cell 27.3, pp. 420–434. issn: 1097-2765. doi: 10.1016/j.molcel.2007.06.016.

Stephens, Matthew (Apr. 1, 2017). “False discovery rates: a new deal”. In: Biostatistics 18.2, pp. 275–294. issn: 1465-4644. doi: 10.1093/biostatistics/kxw041.

Van Nostrand, Eric L. et al. (July 2020). “A large-scale binding and functional map of human RNA-binding proteins”. In: Nature 583.7818. Publisher: Nature Publishing Group, pp. 711–719. issn: 1476-4687. doi: 10.1038/s41586-020-2077-3.

Vehtari, Aki, Andrew Gelman, et al. (June 1, 2021). “Rank-normalization, folding, and localization: An improved $ widehat R $ for assessing convergence of MCMC”. In: Bayesian Analysis 16.2. issn: 1936-0975. doi: 10.1214/20-BA1221. 1903.08008[stat].

Vehtari, Aki, Daniel Simpson, et al. (Mar. 13, 2024). Pareto Smoothed Importance Sampling. doi: 10.48550/arXiv.1507.02646. 1507.02646[stat].

Weyn-Vanhentenryck, Sebastien M. et al. (Mar. 27, 2014). “HITS-CLIP and Integrative Modeling Define the Rbfox Splicing-Regulatory Network Linked to Brain Development and Autism”. In: Cell Reports 6.6, pp. 1139–1152. issn: 2211-1247. doi: 10.1016/j.celrep.2014.02.005.

Youn, Yong Ha et al. (Dec. 9, 2009). “Distinct Dose-Dependent Cortical Neuronal Migration and Neurite Extension Defects in Lis1 and Ndel1 Mutant Mice”. In: The Journal of Neuroscience 29.49, pp. 15520–15530. issn: 0270-6474. doi:10.1523/JNEUROSCI.4630-09.2009.

Zarnack, Kathi et al. (Jan. 31, 2013). “Direct Competition between hnRNP C and U2AF65 Protects the Transcriptome from the Exonization of Alu Elements”. In: Cell 152.3, pp. 453–466. issn: 0092-8674. doi: 10.1016/j.cell.2012.12.023.

Zhang, Chaolin et al. (July 23, 2010). “Integrative modeling defines the Nova splicing-regulatory network and its combinatorial controls”. In: Science (New York, N.Y.) 329.5990, pp. 439–443. issn: 0036-8075. doi: 10.1126/science.1191150.

Zhang, Xiaochang et al. (Aug. 25, 2016). “Cell-Type-Specific Alternative Splicing Governs Cell Fate in the Developing Cerebral Cortex”. In: Cell 166.5, 1147–1162.e15. issn: 0092-8674. doi: 10.1016/j.cell.2016.07.025.

